# The PtdIns3P-binding protein Phafin2 escorts macropinosomes through the cortical actin cytoskeleton

**DOI:** 10.1101/180760

**Authors:** Kay Oliver Schink, Kia Wee Tan, Hélène Spangenberg, Domenica Martorana, Marte Sneeggen, Coen Campsteijn, Camilla Raiborg, Harald Stenmark

**Affiliations:** Centre for Cancer Biomedicine, Faculty of Medicine, University of Oslo, Montebello, N-0379 Oslo, Norway; Department of Molecular Cell Biology, Institute for Cancer Research, Oslo University Hospital, Montebello, 0379 Oslo, Norway; Department of Molecular Microbiology and Genetics, Georg-August-University Göttingen, 37077 Göttingen, Germany; Department of Molecular Medicine, Institute of Basic Medical Sciences, University of Oslo, PO Box 1112 Blindern, 0317 Oslo, Norway

## Abstract

Uptake of large volumes of extracellular fluid by actin-dependent macropinocytosis plays important roles in infection, immunity and cancer development. A key question is how large macropinosomes are able to squeeze through the dense actin network underlying the plasma membrane in order to move towards the cell centre for maturation. Here we show that, immediately after macropinosomes have been sealed off from the plasma membrane, the PH-and FYVE domain-containing protein Phafin2 is recruited by a mechanism that involves binding to phosphatidylinositol 3-phosphate (PtdIns3P) generated in a non-canonical manner. Phafin2 in turn regulates the actin cross-linking protein Filamin A to promote entry of macropinosomes through the subcortical actin matrix and subsequent maturation. Depletion of Phafin2 inhibits macropinocytic internalization and maturation. We conclude that PtdIns3P and its effector Phafin2 are key components of a system that allows nascent macropinosomes to navigate through the dense subcortical actin network.

## Introduction

Macropinocytosis is an endocytosis mechanism that allows cells to take up extracellular fluids and soluble macromolecules by the formation of large vacuoles ^1^. It is important in dendritic cells and macrophages which use this mechanism to sample body fluids for antigens, and for bulk membrane retrieval, e.g. in growth cone collapse during neurite outgrowth. Like many cellular processes, it is subverted and exploited by different pathogens, which either use or even actively trigger macropinocytosis in order to be taken up by host cells ^2^. Macropinocytosis has also recently emerged as a major pathway exploited by cancer cells. Ras-transformed tumor cells, which exhibit high levels of macropinocytosis, utilize this mechanism to take up proteins from the surrounding medium to fuel their increased need for nutrients ^3^. These amino acid-filled macropinosomes can also serve as platforms for mTorc1 signalling. ^4^

Macropinocytosis is an actin-dependent process. Growth factor signalling, active Ras GTPases, or injection of effector proteins by bacterial pathogens trigger actin-driven formation of membrane ruffles. These ruffles can collapse and pinch off from the plasma membrane to form large vesicles filled with extracellular liquid. The mechanism of ruffle closure and pinch-off is yet poorly understood, however, actin-dependent myosin motor proteins have been shown to localize to macropinosome cup and could be involved in closing the ruffle ^5^. How the final scission from the membrane is achieved is unclear – some forms of macropinocytosis appear to require Dynamin, whereas other forms are Dynamin independent. As actin plays a major role in closing the ruffle, it might also be required for scission.

Phosphoinositides are key regulators of macropinocytosis. PtdIns(4,5)P2 and PtdIns(3,4,5)P3 have been shown to localize to macropinosome cups, where they can trigger actin rearrangements by activating different actin-regulating pathways ^6, 7^. After ruffle closure, relatively little is known about the role of phosphoinositides. A phosphatase cascade metabolizing PtdIns(3,4,5)P3 via PtdIns(3,4)P2 and PtdIns3P to PtdIns has been recently described ^8^. It has been proposed that PtdIns3P activates an ion channel, KCa3.1, which is required for macropinosome scission ^8^.

Once macropinosomes are internalized, they gradually acquire endosomal proteins such as the small GTPase Rab5 and its effectors APPL1 and Rabankyrin-5 ^9, 10^. They also acquire stable pools of PtdIns3P, which is synthesized by the PI 3-kinase VPS34. PtdIns3P plays a major role in the transition of APPL1-positive macropinosomes to EEA1-positive macropinosomes ^9^. Macropinosomes follow a similar maturation route as endosomes, sequentially gaining markers of early endosomal, late endosomal and lysosomal identity ^6^.

One of the key questions is how macropinosomes mature immediately after their scission from the plasma membrane and how they gain their endosomal identity. Here, we report a novel maturation stage of macropinosomes immediately after their scission from the plasma membrane and prior to their acquisition of endocytic markers. During this maturation stage, macropinosomes are fully sealed from the plasma membrane but have not yet gained markers of endocytic vesicles such as Rab5 or APPL1. At this stage, their limiting membrane is densely coated with actin, which is stripped from the vesicle during the transition from this earliest maturation step, allowing the macropinosome to escape the cell cortex and acquire endosomal identity. This is associated with a change of macropinosome shape from elongated and multilobed to completely round, the gain of early endocytic markers and the ability to undergo homotypic fusion with other endosomal vesicles. We demonstrate that the PH and FYVE domain-containing protein Phafin2 plays a critical role during this process. Phafin2 is recruited to a transient PtdIns3P pool on newly formed macropinosomes. On these vesicles, it interacts with the actin cross-linking protein Filamin A, and thereby regulates the reorganisation of the actin cytoskeleton to allow the newly formed macropinosomes to shed their actin coat and move towards the cell centre.

## Results

### Phafin2 shows a biphasic localization to macropinosomes

Phafin2 has been reported to reside on endosomes and to regulate – by yet unknown mechanisms – the activity of Rab5 ^11^. Moreover, we have previously shown that Phafin2 is required for effective degradation of endocytosed epidermal growth factor receptors, potentially by regulating endosome fusion ^12^. To further elucidate the mechanism of Phafin2 action on endosomes, we performed live-cell imaging of Phafin2-GFP in retinal pigment epithelial (RPE1) cells stably expressing low levels of the fusion protein. Unexpectedly, we observed two different subcellular localization patterns for Phafin2. In addition to the previously described endosomal localization, we observed a striking localization to large vesicles with a clearly defined lumen. Moreover, in cells with a high number of membrane ruffles, we observed bright, short-lived bursts of Phafin2 localization to vesicles in close proximity to the plasma membrane, in addition to the more stable localization to large vesicles (Figure 1a, movie S1). Dual-colour live cell imaging with plasma membrane tethered mCherry (MyrPalm-mCherry) ^13^ revealed that this transient Phafin2 localization occurred at newly formed macropinosomes, immediately after the formation of new vesicles from cup-shaped membrane ruffles (Figure 1b, c, movie S1).

**Figure 1:**
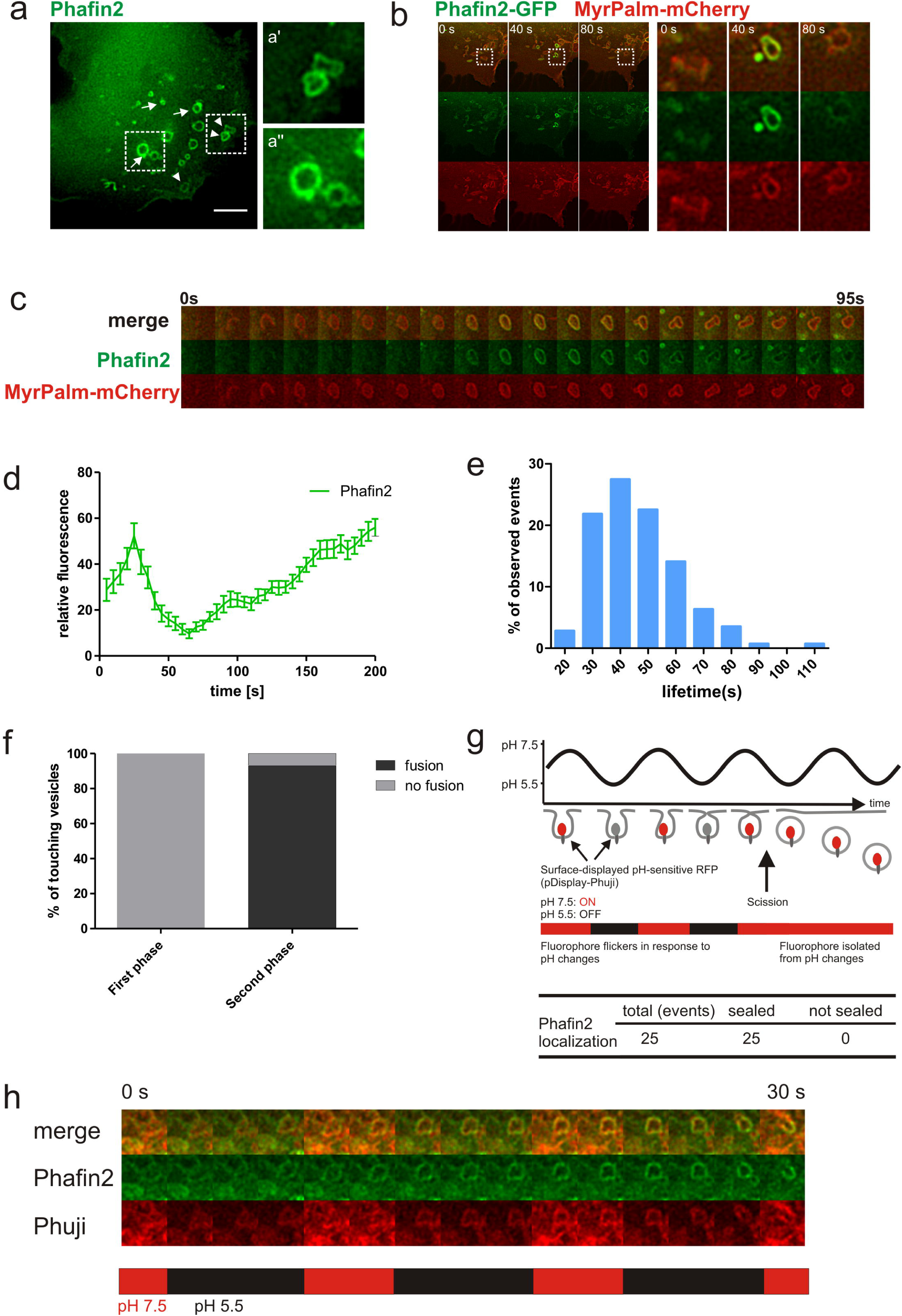
Phafin2 localizes to newly formed macropinosomes. **a)** Micrograph of hTERT-RPE1 cells expressing Phafin2 showing labelling on two different vesicle classes - round, endosome-like vesicles within the cell (arrows, inset a’) and non-uniformly shaped structures close to the cell periphery (arrowheads, inset a’’). Scale bar: 10μm. **b)** Cells co-expressing Phafin2-GFP and MyrPalm-mCherry. Phafin2 transiently localizes to macropinosomes forming from membrane ruffles. Shown are 3 frames from a timelapse sequence, spaced 40s apart. **c)** Sequential images showing Phafin2 dynamics on newly formed macropinosomes; images were acquired every 5s. **d)** Tracking of individual macropinosome shows a biphasic Phafin2 localization to macropinosomes (n=25 macropinosomes, mean + SEM) **e)** Histogram showing the average live-time of the first Phafin2 localization. (n=142 macropinosomes). **f)** Macropinosomes labelled with the first Phafin2 localization are fusion incompetent, whereas vesicles readily undergo fusion during the second phase of localization. (n=53 macropinosomes). **g)** Assay measuring the closure of macropinosomes. Cells displaying pH-sensitive pHuji-RFP on their surface were alternatingly perfused with buffers at pH 5.5 and 7.5 and localization of Phafin2 was evaluated in comparison to Phuji fluorescence (n=25 macropinosomes). **h)** Sequential images of Phafin2 and Phuji expressing cells perfused with pH 5.5 and pH 7.5 imaging buffers, images were acquired every 2s. Note that the displayed sequence shows only a post-scission macropinosome.

Tracking of individual vesicles showed two distinct phases of Phafin2 localization to the same macropinosome, a short-lived (∼40 s), transient localization to structures close to the plasma membrane and a second, long-lasting localization to large vacuoles (Figure 1d,e). The first localization of Phafin2 typically occurred in a single pulse, whereas the second localization was characterized by a gradual increase in fluorescence over several minutes. We also noted that during the first Phafin2 localization, macropinosomes were irregularly shaped and often appeared to be squeezed and subjected to external forces, whereas the second localization occurred on perfectly round macropinosomes (Figure 1a). We noted that macropinosomes during the first phase of Phafin2 localization were unable to undergo homotypic fusion, even if they were in extensive contact with neighbouring vesicles. In comparison, most macropinosomes in the second phase of Phafin2 localization readily fused if they were in contact with other macropinosomes (Figure 1f). Taken together, this indicates that the two phases of Phafin2 localization represent two distinct steps of macropinosome maturation.

### Phafin2 localizes to fully abscised macropinosomes

The observed localization pattern and inability to fuse with neighbouring vesicles allowed for the possibility that the first phase of Phafin2 localization occurred at the forming macropinosome, prior to scission from the plasma membrane. To account for this possibility, we tested whether Phafin2-positive structures were already sealed off from the extracellular environment using a surface-displayed pH-sensitive red fluorescent protein, Phuji ^14, 15^. Cells expressing this marker showed bright surface fluorescence at neutral pH. To assay macropinosome closure, cells were perfused with imaging solutions buffered to pH 5.5 and pH 7.5, alternating every 5 seconds. This pH flickering led to alternating quenching and unquenching of Phuji fluorescence on surface-exposed structures, whereas sealed membranes did not show any fluorescence changes (Figure 1g,h, movie S2). Using this assay, we determined that all observed Phafin2-positive structures were completely sealed (n = 25 macropinosomes).

### Phafin2 labels macropinosomes prior to recruitment of canonical early endocytic markers

To further characterize the spatiotemporal localization of Phafin2, we followed the localization of Phafin2-GFP with different markers of early endosomes and macropinosomes. Individual vesicles were tracked and the relative fluorescence of Phafin2-GFP and the additional marker channel was measured. We first co-expressed Phafin2 with the early endocytic adapter protein APPL1 (Figure 2a,b, movie S3) ^16^. APPL1 has been shown to localize to young macropinosomes prior to their acquisition of EEA1 ^9^. Strikingly, the first phase of Phafin2 localization occurred prior to recruitment of APPL1 to the vesicle membrane. APPL1 only arrived once the first Phafin2 pulse had already dissociated from the vesicle. The second phase of Phafin2 recruitment occurred when levels of APPL1 on the vesicle were already declining. Live-cell imaging of Phafin2 with other markers of early endosomes showed a similar pattern. Imaging of the small GTPases Rab5 (Figure 2e,f) – a marker or early endosomes, Rab31, which has recently been described to localize to early phagosomes (Figure 2g-2f) ^17^, and the Rab5/PtdIns3P effector Rabankyrin-5 (Figure 2c,d), a marker of early macropinosomes ^10^, showed that the first phase of Phafin2 localization occurred prior to the recruitment of these early endosome/macropinosome markers, while the second phase of Phafin2 localization occurred in parallel to recruitment of these markers.

**Figure 2:**
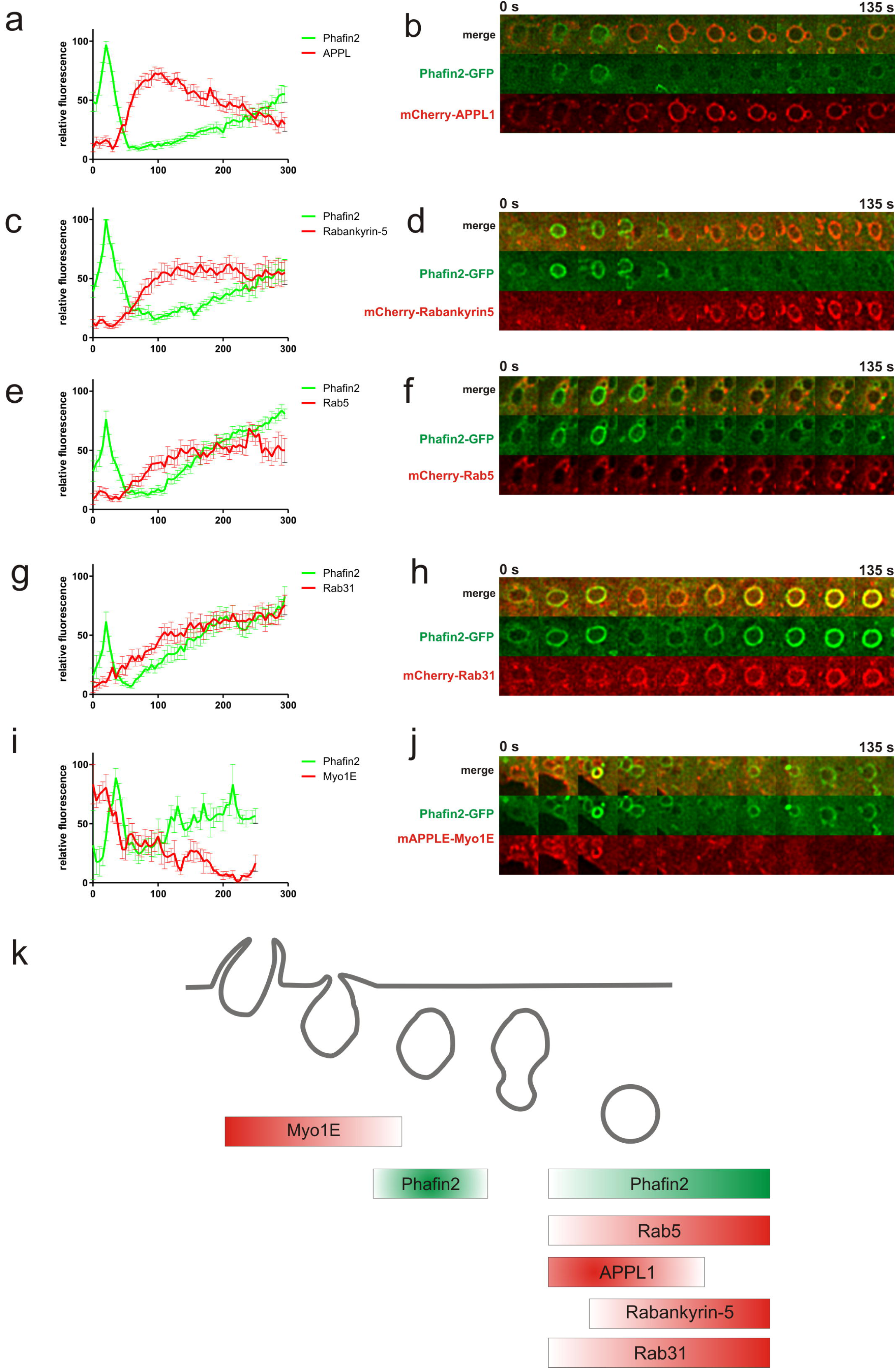
Mapping Phafin2 dynamics in the endocytic pathway. **a)** Phafin2 is recruited to forming macropinosomes prior to APPL1. Forming macropinosomes acquire a brief burst of Phafin2. Following this burst, Phafin2 is completely lost from these vesicles and APPL1 is recruited. During further maturation, APPL1 is gradually replaced by a second, slower recruitment of Phafin2 (n=15 macropinosomes; mean + SEM). **b)** Sequential images showing Phafin2 and APPL1 dynamics on a macropinosome. **c)** Forming macropinosomes acquire a burst of Phafin2, followed by recruitment of Rabankyrin-5. (n=12 macropinosomes, mean + SEM). Macropinosome maturation is accompanied by a second, gradual recruitment of Phafin2. **d)** Sequential images showing Phafin2 and Rabankyrin-5 dynamics on a macropinosome. **e)** Forming macropinosomes acquire a burst of Phafin2, followed by recruitment of Rab5 (n=14 macropinosomes, mean + SEM). Phafin2 shows a second gradual recruitment. **f)** Sequential images showing Phafin2 and Rab5 dynamics on a macropinosome. **g)** Rab31 is recruited after the initial Phafin2 recruitment (n=10 macropinosomes, mean + SEM) **h)** Sequential images showing Phafin2 and Rab31 dynamics on a macropinosome. **i)** Myo1E is recruited earlier than the initial Phafin2 bursts and shows partial temporal overlap during the first phase of Phafin2 recruitment (n=6 macropinosomes, mean + SEM). **j)** Sequential images showing Phafin2 and Myo1E dynamics on a macropinosome. k) Schematic overview of the observed recruitment dynamics.

To further narrow in the timing of the first Phafin2 recruitment, we analysed its localization in relation to the myosin motor protein Myo1E, which is localized to macropinosome cups prior to closure ^5^. The first phase of Phafin2 recruitment occurred after recruitment of Myo1E (Figure 2i,j) and showed only partial temporal overlap in localization, further supporting the finding that the first phase of Phafin2 recruitment occurs directly after plasma membrane scission. Taken together, these data suggest that Phafin2 can label two distinct phases of macropinosome maturation (Figure 2k). Apart from an endosomal localization, Phafin2 recognizes an immediate-early stage of newly-formed macropinosomes that is distinct from and earlier than the better-characterized Rab5-positive maturation step (Figure 2k).

### Phafin2 localization is dependent on its PH and FYVE domains

Next, we asked how these two different macropinosome maturation steps are recognized by Phafin2. Phafin2 is a small protein of 249 amino acids, and its PH and FYVE domains are the main structural features. These potentially phosphoinositide-binding domains are flanked by a short N-terminal predicted α– helical structure which might act as an extension to the PH domain, and a short, highly acidic C-terminal sequence. In order to analyse the role of the different subdomains of Phafin2, we generated a series of truncation mutants lacking different subdomains and assayed their subcellular localization by live cell microscopy (Figure 3a). This analysis showed that both lipid-binding domains play a critical role in localizing Phafin2. The presence of the FYVE domain was required for both the early and late localization of Phafin2 to macropinosomes. In contrast, the PH domain as well as the N-terminal 30 amino acids were critical for localization to the first phase on macropinosomes, but were not required for localization to the second, endosomal phase (Figure 3a). To test whether the first burst of Phafin2 localization to macropinosomes is dependent on the lipid-binding activity of the two domains, we introduced point mutants that abolish lipid-binding activity in the PH and FYVE domains and tested their localization by live cell microscopy. First, we mutated a single amino acid which is critical for PtdIns3P binding in other FYVE domains in the Phafin2 FYVE domain (Phafin2 R176A) ^18^. The R176A point mutation of the FYVE domain abolished membrane association entirely and neither the first nor the second localization to macropinosomes was observed for the point mutant, while wildtype Phafin2 dynamics were unperturbed (Figure 3 b,c; movie S4). Thus, the FYVE domain is essential for the membrane localization of Phafin2.

**Figure 3:**
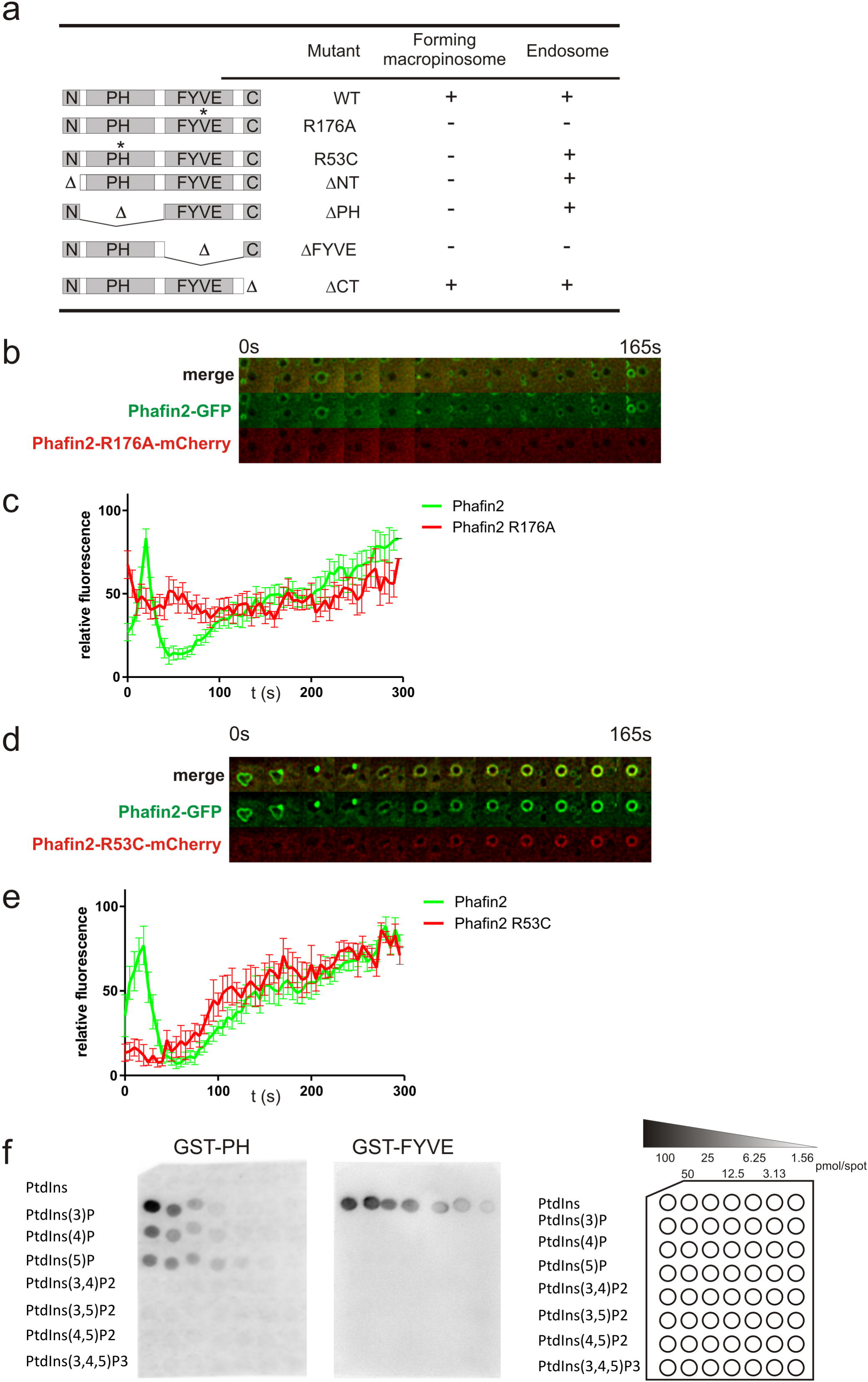
Mutation analysis of Phafin2. **a)** Mutation analysis of Phafin2. Phafin2 truncation and point mutants were tested for their subcellular localization. **b)** Localization dynamics of Phafin2 and Phafin2 with a defective PH domain (R53C) localization (n=6 macropinosomes, mean + SEM). Phafin2 R53C is unable to localize to newly formed macropinosomes, whereas the second, endosomal localization is unaffected. **c)** Sequential images of a macropinosome showing recruitment dynamics of Phafin2 (WT) and Phafin2 (R53C). **d)** Localization dynamics of Phafin2 and Phafin2 with a defective FYVE domain (R176A) (n=15 macropinosomes, mean + SEM). Mutation of the FYVE domain abolishes all membrane localization. **e)** Sequential images of a macropinosome showing recruitment dynamics of Phafin2 (WT) and Phafin2 (R176A). **f)** Protein-lipid binding assay of Phafin2 PH and FYVE domains.

Next, we tested if a functional PH domain is required for Phafin2 localization. To this end, we introduced a mutation into the Phafin2 PH domain, Phafin2 R53C, which has previously been shown to lead to a loss of lipid-binding activity ^19^. The resulting mutant, Phafin2(R53C), still showed the second localization phase to endosomes and macropinosomes, but did not show the first, transient localization to newly-formed macropinosomes (Figure 3d,e; movie S5). This suggests that the PH domain of Phafin2 is not sufficient to mediate membrane association alone and is dispensable for localization to the endosomal stage, but is required to direct Phafin2 to newly-formed macropinosomes. Taken together, these findings indicate that for endosomal localization, the FYVE domain is the main determining factor, whereas the first phase of Phafin2 localization requires a dual recognition module consisting of both PH and FYVE domains.

Based on the observation that Phafin2 labels newly formed vesicles and the strict requirement for both the phosphoinositide-binding PH and FYVE domain, it was tempting to speculate that Phafin2 can recognize a specific phosphoinositide composition on newly-formed macropinosomes. To test this hypothesis, we determined the lipid-binding preference of Phafin2 and analysed Phafin2 localization in relation to different phosphoinositide species. We purified GST-fused PH and FYVE domains expressed in *E.coli* and performed protein-lipid overlay assays ^20^. Both the Phafin2 PH and FYVE domains showed a preference for PtdIns3P. The FYVE domain was highly selective for this phosphoinositide, whereas the PH domain showed a less stringent binding profile and also bound weakly to PtdIns4P and PtdIns5P (Fig 3f) This is in line with previous reports ^19, 21^. As we observed that the N-terminal region of Phafin2 was also required for the first Phafin2 localization phase and its deletion had the same effect as impairing the lipid-binding activity of the PH domain, we asked if this region is required to form a functional PH domain. However, we found that this region was not required for lipid binding nor did its presence change the lipid-binding activity (data not shown).

### Phafin2 recruitment requires PtdIns3P

As we found that both PH and FYVE domains are strictly required for Phafin2 localization and both domains show a preference for PtdIns3P, we reasoned that Phafin2 could recognize different PtdIns3P pools on newly-formed macropinosomes. These two PtdIns3P pools on macropinosomes could be generated by two different pathways – by *de novo* synthesis from PtdIns by VPS34 or PIK3CII isoforms, or as a dephosphorylation product from other 3’ phosphorylated phosphoinositide species. A recent study showed that during the closure of macropinosomes, a cascade of phosphatases catalyses the turnover of PtdIns(3,4,5) via PtdIns(3,4)P2 and PtdIns(3)P to PtdIns (Figure 4a) ^8^. Given the transient nature of the first localization, we reasoned that Phafin2 could be an effector of this phosphatase-generated PtdIns3P pool. To test this hypothesis, we inhibited PtdIns3P *de novo* synthesis using the VPS34 inhibitor SAR405 ^22^. Treatment of Phafin2-GFP expressing cells with SAR405 led to a complete displacement of Phafin2 from the second, endosomal stage (Figure 4b, movie S6), indicating that localization to this stage is PtdIns3P dependent and that VPS34 is the main source of this phosphoinositide. Importantly, SAR-405 treatment did not affect Phafin2 localization to newly-formed macropinosomes. This indicates that the immediate early localization of Phafin2 is independent of VPS34 and further supports the notion that Phafin2 recognizes a phosphatase-generated PtdIns3P pool.

**Figure 4:**
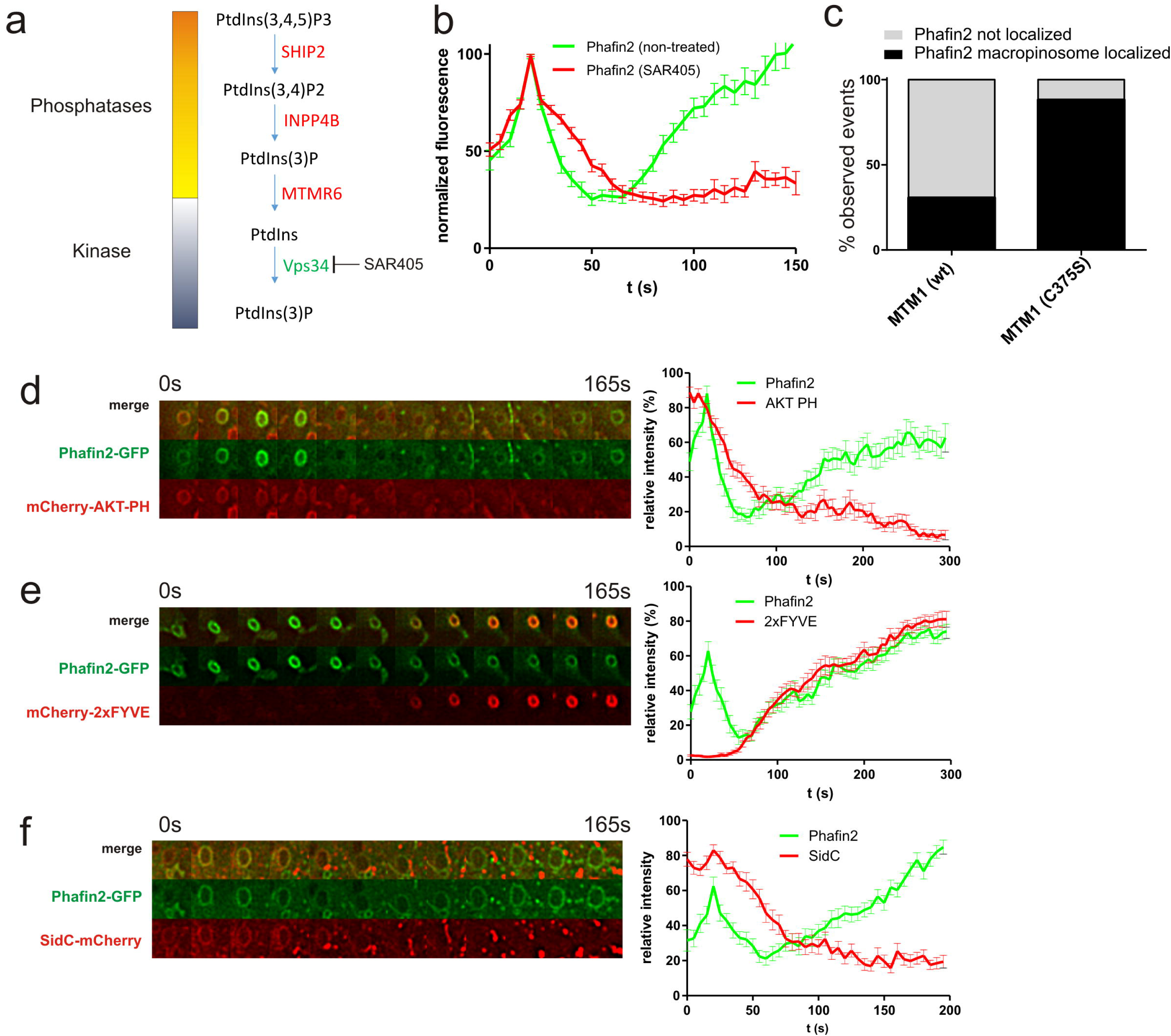
Two independent PtdIns3P pools control the biphasic recruitment of Phafin2. **a)** A phosphatase cascade and the kinase Vps34 generate two independent pools of PtdIns3P on macropinosomes. **b)** Treatment of Phafin2-expressing cells with the Vps34 inhibitor SAR405 does not affect the first localization, but abolishes the second localization phase (n=51 macropinosomes each, mean + SEM) **c)** Overexpression of the PI3 phosphatase MTM1, but not of the catalytically inactive MTM1 (C375S) displaces Phafin2 from macropinosomes (n=75 (wt) and 69 (C375S) cells). **d)** Localization dynamics of Phafin2 in relation to the PtdIns(3,4,5)P3 probe AKT-PH (n=19 macropinosomes, mean + SEM). **e)** Localization dynamics of Phafin2 in relation to the PtdIns3P probe mCherry-2xFYVE (n=20 macropinosomes, mean + SEM). **f)** Localization dynamics of Phafin2 in relation to the PtdIns4P probe SidC (n=25 macropinosomes, mean + SEM)

To test if the first recruitment is PtdIns3P-dependent, we overexpressed the PtdIns3P phosphatase MTM1 and a catalytically inactive mutant (C375S) to indiscriminatingly deplete PtdIns3P from all cellular membranes. Overexpression of wild-type MTM1 led to displacement of Phafin2 from both the first and second localization stage, supporting a role of PtdIns3P for Phafin2 localization (Figure 4c). In contrast, expression of catalytically inactive MTM1 did not affect Phafin2 localization. This indicates that Phafin2 localization is fully dependent on the presence of PtdIns3P and the two distinct phases of Phafin2 localization therefore likely represent two independent PtdIns3P pools that are recognized by Phafin2, one VPS34-derived pool during the endosomal stage and one potentially phosphatase-generated pool at newly formed macropinosomes.

To analyse Phafin2 localization in relation to turnover of PtdIns(3,4,5)P3 to PtdIns3P, we co-expressed Phafin2 together with AKT-PH, a probe for PtdIns(3,4,5)P3 and PtdIns(3,4)P2 ^23^. We found that this probe colocalizes with Phafin2 at newly-formed macropinosomes and that loss of AKT-PH – reporting turn-over of PtdIns(3,4,5)P3 - coincides with the recruitment of Phafin2 (Figure 4d). In order to visualize PtdIns3P on newly-formed macropinosomes, we also analysed the localization of Phafin2 in relation to mCherry-2xFYVE, a probe specific for PtdIns3P ^24^. Surprisingly, we did not observe any colocalization between Phafin2 and the 2xFYVE probe at newly-formed macropinosomes, whereas we observed a nearly perfect colocalization on endosomes and fully-formed macropinosomes (Figure 4e). mCherry-2xFYVE was not present on newly-formed macropinosomes or membrane ruffles and could only be detected on later, endosomal vesicles together with the arrival of early endocytic markers. This might be due to low levels of PtdIns3P on the plasma membrane, which could require additional cues, e.g. by coincidence sensing of PtdIns3P by both the PH and FYVE domain or interaction with protein partners.

As the Phafin2 PH domain does not only bind to PtdIns3P, but also to PtdIns4P, we could not exclude that the first localization of Phafin2 might rely on coincidence sensing of transiently generated PtdIns3P by the FYVE domain and plasma-membrane-derived PtdIns4P by the PH domain instead of PtdIns3P alone. Co-expression of Phafin2 with SidC, a high-affinity probe for PtdIns4P ^25^ showed colocalization of Phafin2 with SidC during the first Phafin2 localization (Figure 4f). We reasoned that if Phafin2 requires PtdIns4P, overexpression of SidC should outcompete Phafin2. However, we found that even cells expressing high levels of SidC-mCherry readily recruited Phafin2 to newly-formed macropinosomes and did not differ from control cells expressing only mCherry (not shown). Furthermore, we found no significant correlation between the intensities of Phafin2 and SidC during the first peak of Phafin2 recruitment (Spearman’s rho = −0.059, p=0.84). Based on these observations, it is unlikely that Phafin2 requires PtdIns4P for its recruitment.

### Absence of Phafin2 causes defects in macropinocytosis

Based on the localization of Phafin2 to macropinosomes, we asked if Phafin2 has a functional role in macropinocytosis. To test whether cells lacking Phafin2 are affected in macropinocytosis, we first assayed for defects in fluid-phase uptake. To this end, we depleted Phafin2 using siRNA and measured uptake of fluorescently labelled dextran using flow cytometry. We found that uptake of dextran was significantly reduced by 40 - 50 % if Phafin2 was depleted (n = 3 experiments, two siRNAs, >8000 cells per condition) (Figure 5a).

**Figure 5:**
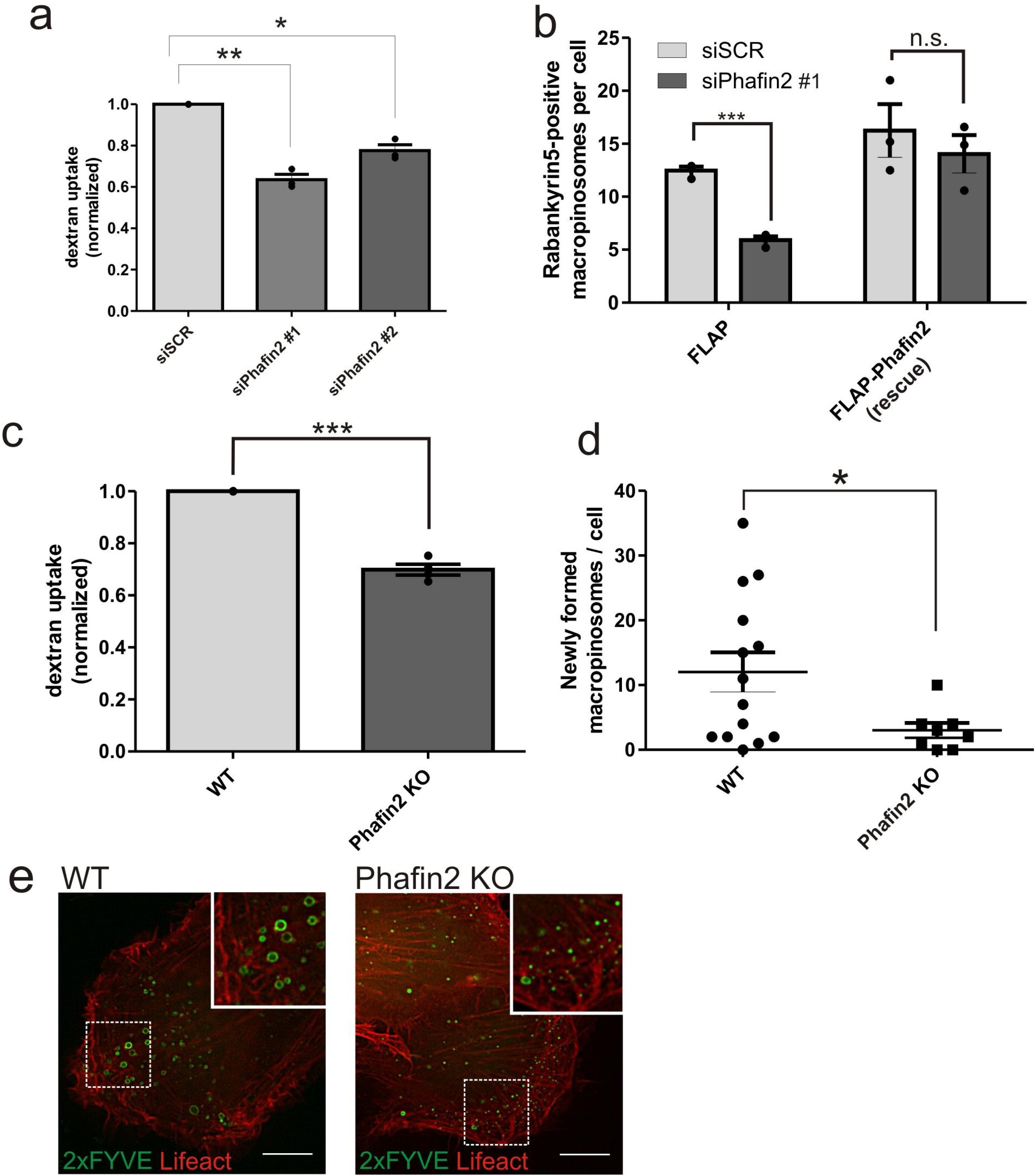
Phafin2 is required for efficient macropinocytosis. **a)** Measurement of dextran uptake (70 kDa) in Phafin2 depleted cells (3 experiments, mean + SEM, > 8000 cells per experiment and condition, p=0.0051 and p=0.0161). **b)** Formation of Rabankyrin-5 positive macropinosomes in Phafin2 depleted cells and rescue with siRNA resistant Phafin2 (3 experiments , mean + SEM , > 2500 cells per condition, p= 0.0002 (Phafin2 depletion) and p=0.451 (rescue)). **c)** Measurement of dextran uptake (10kDa) in Phafin2 knockout cells (mean of 4 experiments + SEM, > 2000 cells per experiment and condition, p=0.0007). **d)** Formation of new macropinosomes (> 1 um in diameter) in Phafin2 wild-type and KO cells. Macropinosome formation was scored over 20 min of imaging time. (n=14 (wt) and 8 (KO) cells, mean + SEM, p=0.043). e) Exemplary image showing Macropinocytosis in wild-type and Phafin2 knockout cells (single frame extracted from the movies analyzed in (d)).

As we had established that Phafin2 arrives at a very early phase of macropinosome maturation, but after scission from the membrane, we asked why depletion of Phafin2 would result in less fluid phase uptake. The protein Rabankyrin-5 has been shown to label newly formed macropinosomes and we found that Rabankyrin-5 arrives shortly after the initial Phafin2 localization ^10^. Therefore, we reasoned that a Phafin2-dependent macropinocytosis effect could affect maturation of newly-formed macropinosomes to the Rabanykrin-5 positive stage. To test this hypothesis, we labelled cells depleted for Phafin2 for Rabankyrin-5 and automatically scored for large Rabankyrin-5 positive structures. Using this assay, we found that Phafin2 depletion severely reduced the number of Rabankyrin-5 labelled macropinosomes (Figure 5b, Figure S1). Stable expression of siRNA-resistant Phafin2 could rescue this defect, indicating that the observed effects of the siRNA were specific. We observed that cells depleted for Phafin2 consistently showed only very small Rabankyrin5-labelled endosomes, whereas cells expressing sRNA-resistant Phafin2 did show large macropinosomes (Fig. 5b, Figure S1).

To further corroborate the role of Phafin2 in macropinocytosis, we generated RPE1 cells completely lacking Phafin2 using CRISPR/Cas9-mediated genome editing (Figure S2).

We first tested whether these Phafin2 -/- cells show similar uptake defects as cells depleted for Phafin2 using siRNA. To this end, we assayed fluid-phase uptake using flow cytometry. Similar to cells depleted for Phafin2 by siRNA, Phafin2 KO cells showed reduced dextran uptake to approximately 70 % of wild-type conditions (Figure 5c).

We then used the Phafin2 -/- cell line to follow macropinocytosis in these cells by live cell imaging. Cells were transfected with GFP-2xFYVE to label endosomal vesicles and with Lifeact-SNAP to label membrane ruffles. We selected ruffling cells and followed the formation of new macropinosomes using GFP-2xFYVE. In wild-type cells, we readily observed the formation of multiple large vesicles (diameter > 1 μm). These vesicles were not formed by homotypic fusion, but nearly exclusively by de-novo by macropinocytosis (Fig 5d, e and movie S7). In contrast, despite heavy ruffling in Phafin2 KO cells, we only occasionally observed the formation of large vesicles by macropinocytosis (Fig. 5d,e and movie S6). Overall, 2xFYVE-labelled vesicles were much smaller in KO cells and if ruffling resulted in successful macropinocytosis, the resulting macropinosomes were consistently smaller than in wild-type cells. These results further establish a role of Phafin2 in early steps of macropinocytosis (Fig. 5d, e).

### Phafin2 interacts with the actin crosslinking protein Filamin A

As Phafin2 does not have any obvious catalytic or regulatory domains apart from its PH and FYVE domain, we reasoned that it is likely an interaction partner of Phafin2 that is required for its function during macropinocytosis. In order to investigate the molecular function of Phafin2 and to identify this interaction partner, we screened for interactors of Phafin2 by means of a yeast two-hybrid screen, as well as by affinity purification of GFP-tagged Phafin2 followed by mass spectrometry.

Using both methods, we identified the actin crosslinking protein Filamin A as potential interaction partner of Phafin2. Filamin A was found to be 7-fold enriched in mass spectrometry, and we identified two independent fragments coding for an overlapping region of Filamin A in a yeast two-hybrid screen.

Filamin A contains two N-terminal Calponin-homology (CH) domains, followed by 24 IG repeats (Figure 6a) ^26^. The CH domains allow Filamin to bind to actin filaments, whereas the repeats form an extended rod-shaped structure. Homodimerization at the C-terminal regions allows Filamin A to adapt a V-shaped structure which enables crosslinking of actin filaments to a dense network. We identified a fragment coding for amino acids 186 – 368 of Filamin A which interacted with Phafin2 by yeast two-hybrid assays; this region encompasses the second CH domain and the first IG repeat (Figure 6a). To verify the interaction of Filamin A with Phafin2, we measured the interaction using a yeast two-hybrid β-galactosidase assay. This assay further supported the interaction of Phafin2 with Filamin A (Figure 6b). In order to identify the region of Phafin2 interacting with Filamin A, we performed two-hybrid interaction assays with truncation mutants of Phafin2. This revealed that the interaction surface of Phafin2 with Filamin A is localized in the N-terminal region of Phafin2 including the PH domain, as truncation mutants lacking the N-terminus or the PH domain were unable to interact in two hybrid assays. In comparison, mutants lacking the FYVE domain or the C-terminus showed robust interaction (Figure 6b). From this analysis, we conclude that the N-terminus of Phafin2 interacts with the actin-binding domain (ABD) of Filamin A.

**Figure 6:**
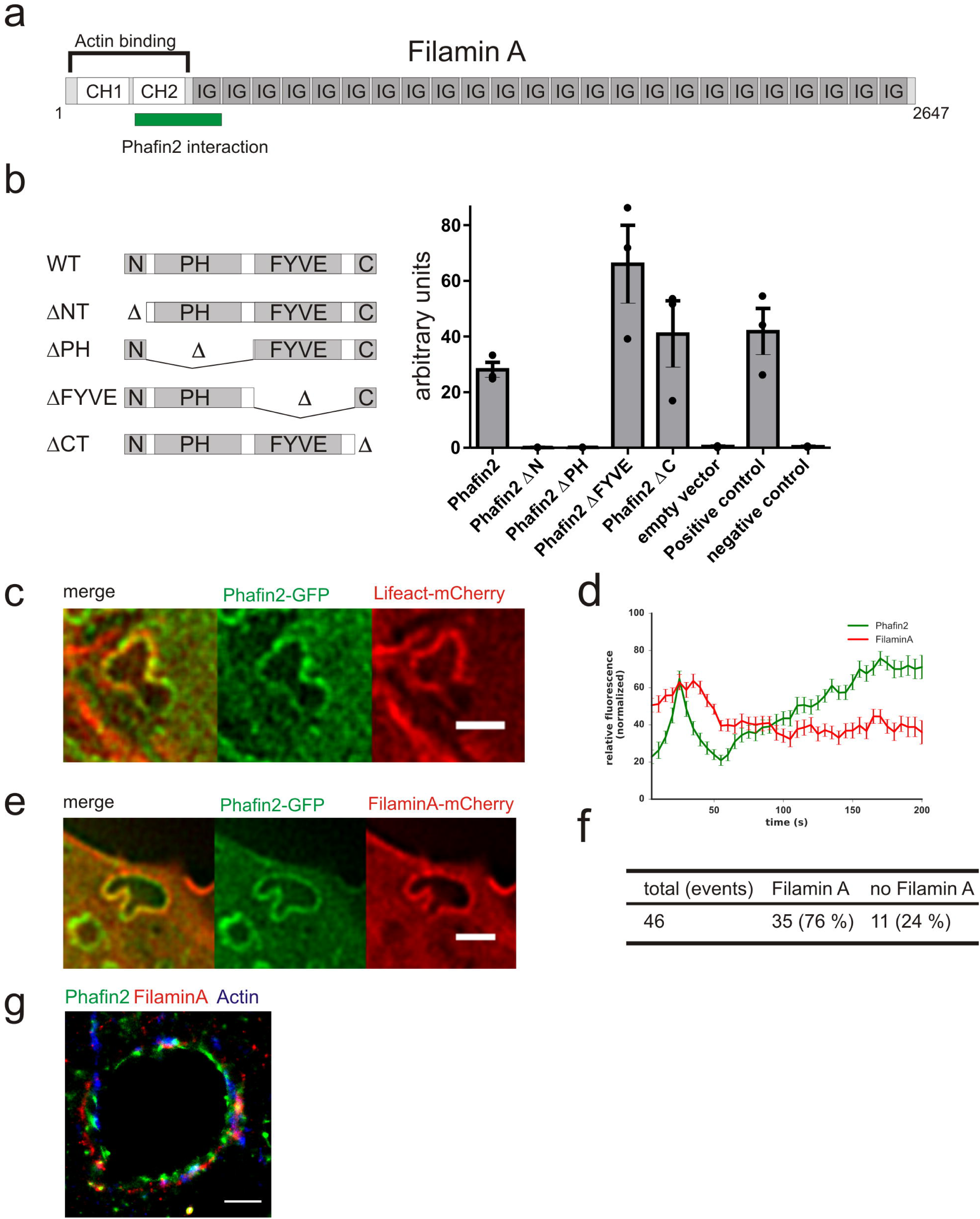
Phafin2 interacts with the actin crosslinking protein Filamin A. **a)** Filamin A domain structure and the identified Phafin2 binding site. **b)** Two-hybrid analysis of Phafin2 truncation mutants with Filamin A and mapping of the Phafin2 domains binding to Filamin A. **c)** Actin dynamics on Phafin2-labelled macropinosomes. Single frame extracted from movie S7 (scalebar = 2μm). **d)** Localization dynamics of Phafin2 and Filamin A on macropinosomes (n = 46 macropinosomes). **e)** Colocalization of Phafin2 with Filamin A-mCherry. Filamin A-mCherry forms a coat around newly-formed macropinosomes. Single frame extracted from movie S12. (scalebar = 2μm). **f)** Quantification of Filamin A association with Phafin2-positive macropinosomes (n = 46 forming macropinosomes). **g)** 3-color dSTORM image showing Phafin2, FilaminA and Actin coating a macropinosome (scalebar = 1μm).

### The first phase of Phafin2 localization coincides with actin rearrangements around macropinosomes

Based on the effects of Phafin2 depletion on the structure of the actin cytoskeleton and the identification of Filamin A as Phafin2 interaction partner, we asked how the actin cytoskeleton is arranged during Phafin2 localization to macropinosomes and whether Filamin A is associated with Phafin2-labelled macropinosomes. As we observed that newly formed Phafin2-labelled macropinosomes appeared to be subject to mechanical forces, we reasoned that there might be a close relationship between these vesicles and the actin cytoskeleton. Live-cell imaging of Phafin2 together with Lifeact-mCherry showed that newly-formed Phafin2-labelled macropinosomes are coated with a dense network of actin filaments (Figure 6c, movie S8). The first recruitment of Phafin2 coincided with a restructuring of this actin network. Actin-coated macropinosomes were subject to squeezing forces that pushed them into the inner parts of the cell (Figure 6c, movie S7). Ultimately, we observed that gaps in the actin cytoskeleton lining the macropinosomes were formed which allowed them to leave their actin coat behind and acquire a completely round-vesicle-like shape (movie S7). A key to understanding macropinosome maturation is thus to characterize how these gaps in the subcortical actin network are formed.

We therefore asked whether the Phafin2 interacting protein Filamin A is localized to Phafin2-decorated macropinosomes and could be binding to Phafin2 to organize the observed changes of the actin coat surrounding newly formed macropinosomes. We observed that newly formed macropinosomes were not only coated with actin, but also with Filamin A (Figure 6e, movie S9). The majority of newly formed macropinosomes gained a transient Filamin A coat (76% of newly formed macropinosomes, n = 46 macropinosomes (Figure 6f)). Three-color dSTORM revealed a close association of both Actin and Filamin A with Phafin2 coated vesicles (Figure 6g). Dissociation of Filamin A coincided with recruitment and dissociation of Phafin2 (Figure 6d, movie S9). The observed rearrangement of the actin/Filamin A cytoskeleton consistently coincided with loss of the first Phafin2 localization and the transition to the round vesicular shape, indicating that this is a critical step in the maturation of macropinosomes (Figure 6e, movie S9). Likewise, we observed that newly-formed macropinosomes entered the cell through a gap in the surrounding actin and Filamin A coat, indicating that at these places, actin depolymerization allows the vesicle to enter the cell (Figure 6e, movie S10).

Next, we asked whether deletion of Phafin2 affects the behaviour of Filamin A at forming macropinosomes. To this end, we transfected both wild-type and Phafin2 -/- cells with Filamin A and followed the formation of macropinosomes in these cells. In both cases, we observed localization of Filamin A to membrane ruffles and newly-formed macropinosomes, suggesting that Phafin2 does not completely block initial macropinosome formation and is not required to recruit Filamin A (Figure 7c,d). However, we observed that in Phafin2 KO cells, the majority of macropinosomes did not enter further into the cell, but collapsed directly after their formation and appeared to fuse back to the plasma membrane (Figure 7a, d). Closer inspection of these events revealed that the majority of the collapsing macropinosomes retained Filamin A at their limiting membrane (Figure 7b, d; Figure S3B; movie S11). In comparison to this, surviving macropinosomes consistently lost their Filamin A coat (Figure 7b, c). The fraction of surviving macropinosomes in wild-type cells was significantly higher than in Phafin2 -/- cells (Figure 7b), and we observed that essentially all surviving macropinosomes lost Filamin A from their limiting membrane (Figure 7b, c; Figure S3a; movie S11). In cases where macropinosomes did not completely remove Filamin A in wild-type cells, these macropinosomes were visibly subjected to external forces and frequently collapsed (Figure 7b). Taken together, we observed a strong correlation between the survival of macropinosomes and the loss of Filamin A from their limiting membrane (Pearson’s Rho = 0.8126, p < 0.0001; 50 events; data from both WT and Phafin2 -/- cells). This suggests that shedding of Filamin A from the macropinosome membrane is a critical step of macropinosome maturation. Our results also suggest that the observed macropinocytosis defect in response to Phafin2 depletion or deletion is not directly linked to defects in macropinosome formation or scission, but rather due to a maturation defect that could result in back-fusion of the macropinosome with the plasma membrane, thereby resulting in fewer macropinosomes entering the cell.

**Figure 7:**
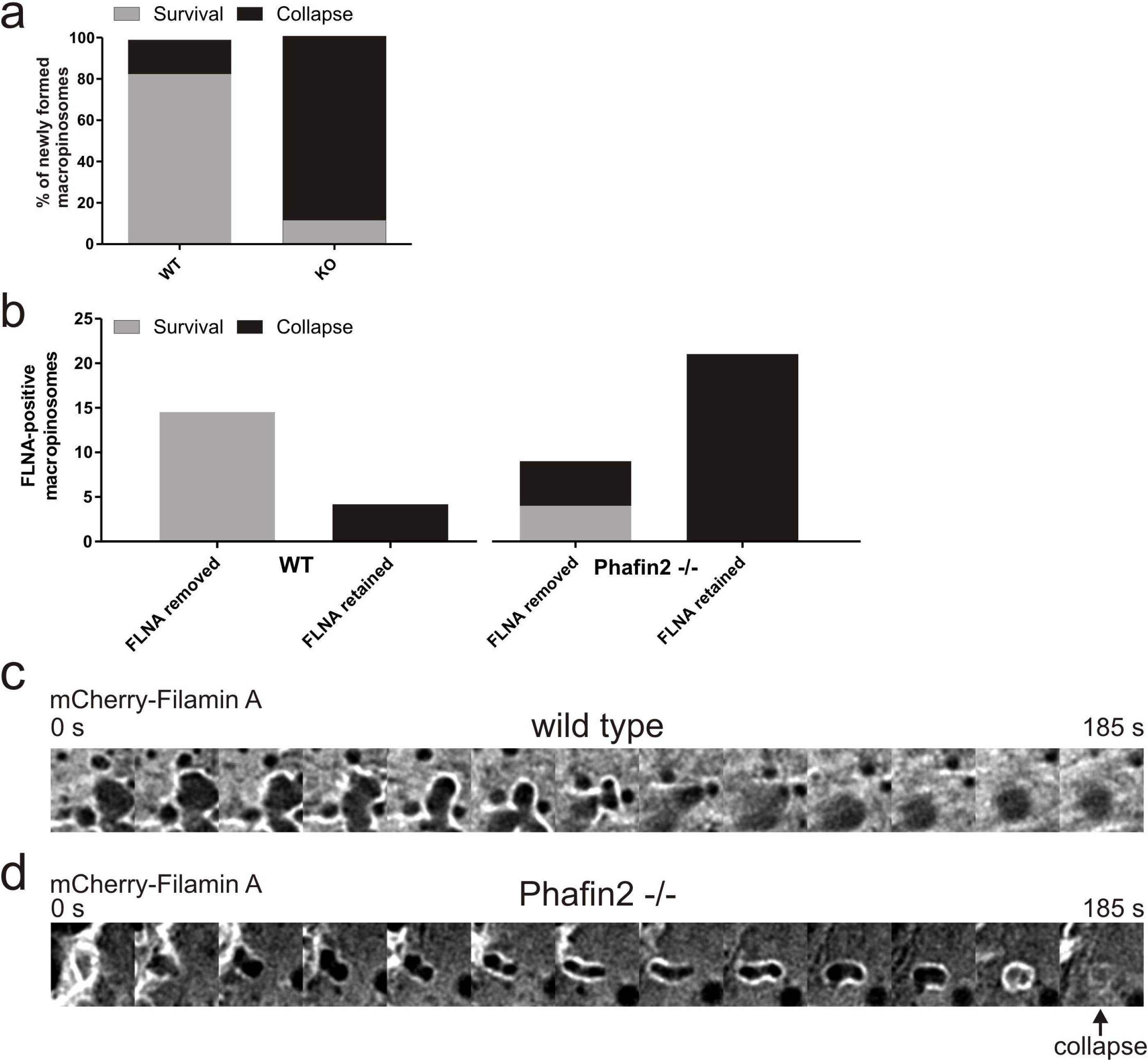
Phafin2 mediates shedding of Filamin A from newly formed macropinosomes. **a)** Analysis of macropinosome dynamics and survival in wild-type and Phafin2 -/- cells. Forming macropinosomes were scored for their transition to round macropinosomes or back-fusion to the plasma membrane (data from 18 (wt) and 34 (KO) macropinosomes). **b)** Scoring the relation between Filamin A shedding and macropinosome survival in wild-type and Phafin2 KO cells. Macropinosome tracks from (a) were scored for shedding of Filamin A and their transition to later macropinosome stages. (n=18 (wt) and 34 (KO) macropinosomes). **c)** Image series showing shedding of Filamin A from newly-formed macropinosomes and their transition to later macropinosomes in wild-type cells. **d)** Image series showing failed shedding of Filamin A from newly-formed macropinosomes in Phafin2 KO cells and the of the macropinosome.

## Discussion

In this study, we provide new insight into the mechanisms of early macropinocytosis and its regulation by PtdIns3P. We demonstrate a new maturation stage of macropinosomes prior to their acquisition of markers of early endosomal identity. After scission from the plasma membrane, newly formed macropinosomes gain PtdIns3P by a VPS34-independent mechanism, which is recognized by Phafin2. This maturation stage is distinct from endosomal stages. Depletion of Phafin2 impairs macropinocytosis and Phafin2 involved in coordinating actin dynamics on newly formed macropinosomes. After scission from the membrane, newly formed vesicles are coated by a dense actin meshwork crosslinked by Filamin A. Gaps in this network allow the macropinosome to escape its actin coat, thereby enabling maturation and progression into the endo-lysosomal pathway. We conclude that Phafin2 interacts with and inactivates Filamin A and thereby allows newly formed vesicles to remove their actin coat.

Phosphatase-produced PtdIns3P was proposed to be a critical factor for completion of macropinocytosis by regulating the KCa3.1 ion channel ^8^. We propose that Phafin2 is a novel effector of this transiently generated PtdIns3P pool. Phafin2 binds to PtdIns3P with both its PH and FYVE domains and both domains are required for recruitment of the protein to newly-formed macropinosomes, supporting the finding that a transient PtdIns3P pool is generated on nascent macropinosomes ^8^. This is further supported by our observation that phosphatase-driven depletion of PtdIns3P displaces Phafin2 from both newly-formed and mature macropinosomes. Finally, we show that this PtdIns3P pool is formed independently of VPS34 activity, consistent with the notion that PtdIns3P in this context is generated by metabolizing other 3’-phosphorylated PI species ^8^. Taken together, this strongly suggests that Phafin2 can sense a transient, VPS34-independent PtdIns3P pool on nascent macropinosomes.

Phafin2 recruitment occurs in parallel to the turnover of other PI species, such as PtdIns(4,5)P2 and PtdIns4P, suggesting that directly after being separated from the plasma membrane, macropinosomes undergo massive changes of their PI content. This could drive maturation of the macropinosome by stripping it of its plasma membrane identity. While it has been shown that ruffles limit membrane diffusion of larger molecules ^27^, it stands to reason that PIs as small, mobile molecules would not be strongly affected by these diffusion barriers. Therefore, prior to scission, PIs would be able to diffuse in and out of the macropinosome cup and equilibrate with the surrounding plasma membrane. However, directly after scission, the available pool of PIs on the limiting membrane of the macropinosome would be constant and subject only to turnover by kinases and phosphatases.

The transient and specific localization of Phafin2 to newly formed macropinosomes suggests a potential function during the maturation or trafficking of these vesicles. Cells lacking Phafin2 showed reduced fluid phase uptake and fewer macropinosomes. While this phenotype could result from defects in macropinosome formation or scission, we demonstrate that Phafin2 arrives after scission from the plasma membrane is complete and therefore its absence should not directly affect macropinosome formation. Consistent with this notion, we observed that cells deleted for Phafin2 were still ruffling and forming cup-shaped ruffles.

We observed that Phafin2 interacts with the actin-crosslinking protein Filamin A and these two proteins transiently co-localize on nascent macropinosomes directly after their formation. This localization coincides with actin reorganization on the macropinosome membrane. Dissociation of Filamin A from the limiting membrane strongly correlates with survival and successful maturation of the macropinosome to the endosomal stage.

The identified interaction region of Filamin A with Phafin2 encompasses part of the Filamin A ABD, suggesting a potential mechanism for the observed actin rearrangements. Binding of proteins, e.g. Calmodulin, to the Filamin A ABD has been shown to displace Filamin A from the actin cytoskeleton ^28^. Binding of Phafin2 to the ABD could weaken the Filamin A interaction with actin, thereby weakening actin crosslinking. This would allow actin reorganizing proteins, e.g. actin capping and bundling proteins, to open and stabilize gaps in the actin cage and thus allow the macropinosome to shed its actin coat and move towards the cell centre.

Our collective findings suggest the following model (Figure 8). Formation of macropinosomes requires actin-driven membrane ruffling and myosin-controlled constriction of the macropinosome cup. Macropinosome cups are enriched in PtdIns(3,4,5)P3 and PtdIns(4,5)P2, and thus serve as potent nucleation platforms for actin, which ultimately drives the formation of vesicles ^7, 29^. At the same time, this enrichment in actin presents a problem, as scission from the plasma membrane results in a vesicle which is caged in actin and trapped in close proximity to the plasma membrane with its actin nucleating factors. In order to move into the cell and enter the endocytic pathway, it would have to remove actin nucleating factors from its limiting membrane and shed its actin coat. Dephosphorylation of PtdIns(3,4,5)P3 and PtdIns(4,5)P2 would result in dissociation of actin regulators, such as GEFs for Rho GTPases, while Rho GAPs could be recruited by one of the dephosphorylation products to inactivate Rho GTPases and terminate actin polymerization. Indeed, the uptake of large particles by phagocytosis requires inactivation of Rho GTPases by phosphoinositide-driven recruitment of Rho-GAPs ^30^. Simultaneously, transient production of PtdIns3P from these dephosphorylation cascades would result in transient recruitment of high levels of Phafin2, which – by interacting with Filamin A and locally competing for its binding to actin - could lead to a zone of weekly crosslinked actin filaments around the macropinosome. Since the area closest to the plasma membrane will have the densest actin network, whereas macropinosomes will only have a thin lining of actin, localized depolymerization, together with contractile forces of the cell cortex, will ultimately push the vesicle out of the region with dense cortical actin and into the inner volume of the cell, thereby allowing its maturation.

**Figure 8:**
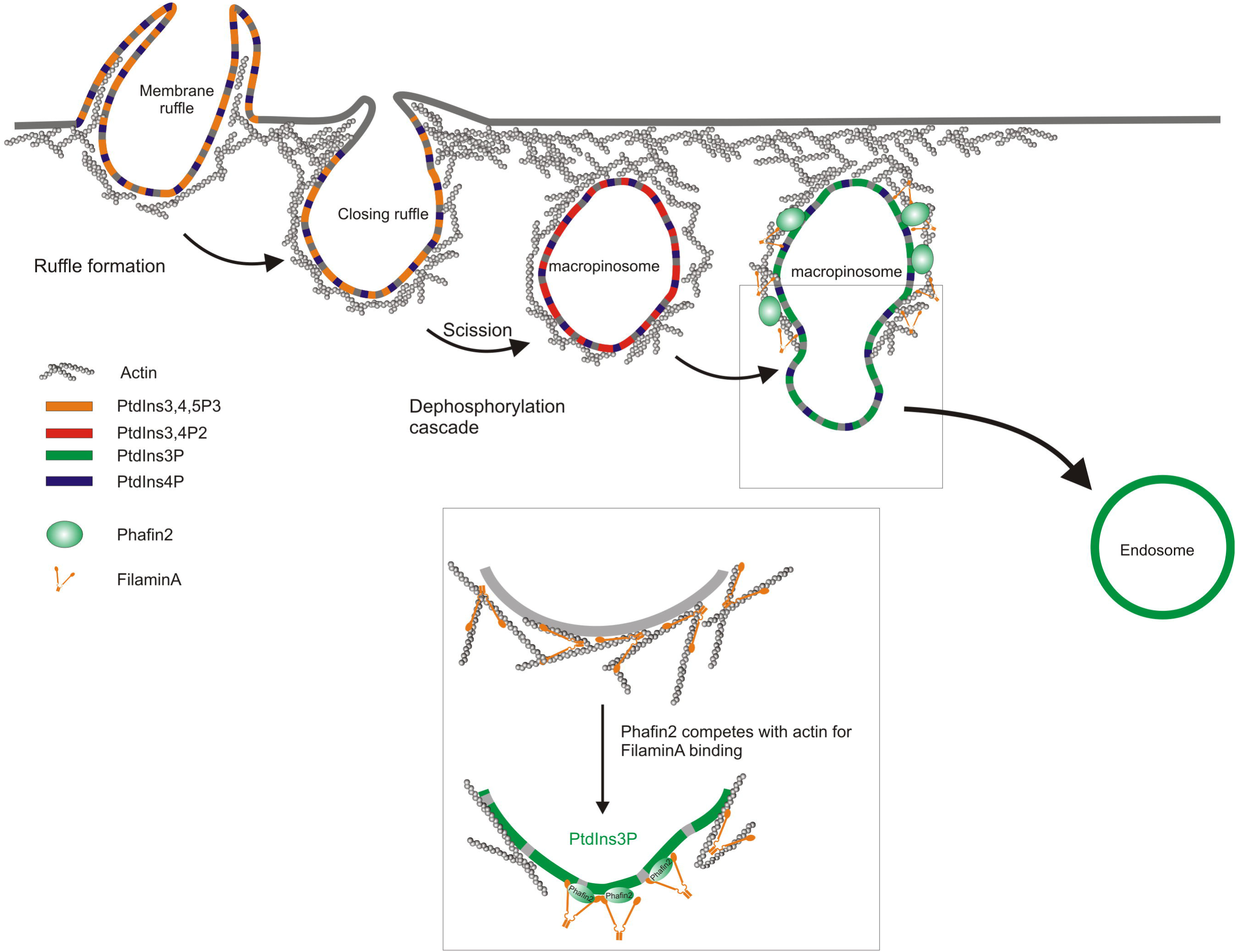
Model of Phafin2-mediated macropinosome maturation. Macropinosomes arise from membrane ruffles and pinch off. Newly formed macropinosomes are enveloped in a dense actin meshwork. Phosphoinositide turnover generates a transient PtdIns3P pool that is recognized by Phafin2. This leads to transient, high levels of Phafin2 at the limiting membrane of the macropinosome. Phafin2 can bind to the actin-binding domain of Filamin A, thereby competing with actin for Filamin A-binding and locally lowering the degree of crosslinking, thereby allowing the macropinosome to escape from the surrounding actin coat.

We note that several observations reported on the bacterial invasion of mammalian cells are in concordance with our proposed model and support the notion that nascent macropinosomes transitioning to the endosomal stage are enriched in machinery to remodel dense actin coats. The intracellular bacterial pathogen *Shigella flexneri* gains entry into the cell by inducing extensive actin remodelling and membrane ruffling, but is then enclosed in an actin-dense meshwork surrounding the plasma membrane-derived vacuole ^31^. Disassembling this coat and subsequent vacuole rupture requires close contact between newly formed macropinosomes and the vacuole, without which the bacterial vacuole remains trapped in an actin cage ^31^. In future studies, it will be interesting to learn whether the Phafin2-Filamin A axis is target for microbial factors that subvert macropinocytosis.

## Author contributions

K.O.S and H.St. conceived and supervised the study. K.O.S. designed experiments, generated constructs, lentiviruses and stable cell lines, performed CRISPR/Cas9 knockouts, performed live cell and superresolution microscopy, flow cytometry, wrote image and data processing software and analyzed data. K.W.T. generated constructs, lentiviruses and cell lines, performed live cell imaging and quantified data, H.Sp. performed image quantification and RNAi experiments. D.M. performed flow cytometry and protein-lipid interaction assays, generated constructs and cell lines. M.S. performed GFP trap experiments. C.C. generated constructs and cell lines and performed GFP trap and proteomics analysis. C.R. performed high content imaging and analysis. K.O.S. and H.St wrote the manuscript with input from all co-authors

## Acknowledgements

We thank Dr. Frauke Ackermann for critical reading of the manuscript and useful discussions and Eva Rønning for technical support with yeast two-hybrid screening. The Advanced Light microscopy core facility at Oslo University Hospital is acknowledged for providing access to relevant microscopes. K.O.S is a research fellow and CR a senior research fellow of the Norwegian Cancer Society. C.C. is a young investigator of the Research Council of Norway. This work was partly supported by the Research Council of Norway through its Centres of Excellence funding scheme, project number 179571.

## Methods

### Cell lines

All experiments were performed in hTert-RPE1 cells (ATCC CRL-4000). H2B-mCherry expressing cells were generated by transfection using pH2B-mCherrry-IRES-Neo, Neomycin selection and picking of single clones. Other stable cell lines were lentivirus-generated pools. Third-generation lentivirus was generated using procedures and plasmids as previously described ^32^. Briefly, tagged fusions of transgenes were generated as Gateway ENTRY plasmids using standard molecular biology techniques. From these vectors, lentiviral transfer vectors were generated by Gateway LR recombination into lentiviral destination vectors (Gateway-enabled vectors derived from pCDH-EF1a-MCS-IRES-PURO (SystemBiosciences)). VSV-G pseudotyped lentiviral particles were packaged using a third-generation packaging system (Addgene plasmids 12251, 12253, 12259). Cells were then transduced with low virus titers (multiplicity of infection < 1) and stable expressing populations were generated by antibiotic selection. Detailed cloning procedures are available from the authors.

### Antibodies

The following primary antibodies were used: Phafin2 antibody, Sigma-Aldrich (HPA024829). Filamin A antibody, Abcam (ab76289). Atto-488 conjugated anti-GFP nanobody, Chromotek (gba488). γ-Tubulin antibody, Sigma-Aldrich (T5326), aTubulin antibody, Sigma-Aldrich (T5168), anti-Rabankyrin-5 antibody (Abnova, H0005 1479-B01P).

### Plasmids

pEYFP-Rabankyrin-5 was a gift from Marino Zerial ^10^; EYFP was exchanged with mCherry. The following plasmids were obtained from Addgene:

Myo1E-pmAppleC1 was a gift from Christien Merrifield (Addgene #27698, ^15^), pH2B-mCherrry-IRES-Neo was a gift from Daniel Gerlich (Addgene #21044, ^33^), pDisplay-Phuji was a gift from Robert Campbell (Addgene #61556, ^14^), pEGFP-AKT-PH was a gift from Tobias Meyer (Addgene #21218, ^34^); EGFP was replaced with mCherry. pmCherry-Filamin A was a gift from Michael Davidson (Addgene plasmid # 55047), pLifeact-mTurquoise2 was a gift from Dorus Gadella (Addgene plasmid #36201, ^35^; mTurquoise2 was replaced with mCherry and SNAP. pX458 was a gift from Feng Zhang (Addgene plasmid # 48138, ^36^)

All other plasmids were generated using standard cloning procedures and are listed in Supplemental table S1.

### Transient transfection of cells

hTert-RPE1 cells were transfected with Fugene 6 using a ratio of 3:1 of reagent to DNA. In most cases cells were transfected in Mattec 3.5 cm dishes using 6 ul of Fugene 6 and 2 ug of DNA. Medium was replaced 12 h after transfection to remove transfection reagent, and cells were imaged between 16h and 20h after transfection.

### CRISPR/Cas9-mediated deletion of Phafin2

hTert-RPE1 cells deleted for Phafin2 were generated using CRISPR/Cas9. Guide RNAs were designed using Benchling software (http://www.benchling.com). For deletion of Phafin2, a guide RNA binding in directly at the start codon in Exon2 and one binding after Exon2 was chosen (gRNA1: 5’-AGGCTATTAGTGAAAGATGG-3’; gRNA2: 5’-TGGGAGTATTTAATCAGGTG-3’). As the whole ORF of Phafin2 is contained in this exon, we reasoned that the whole ORF should be excised. pX458-derived plasmids encoding both Cas9-2A-GFP and the respective gRNA were transfected using Fugene6. 48 h post-transfection, GFP-positive cells were sorted and seeded out in several dilutions to obtain single colonies, which were picked and characterized. Clones lacking Phafin2 were identified by western blotting, and the introduced mutations were characterized by PCR followed by cloning and sequencing. We established a Phafin2 clone lacking the whole Exon 2 in one allele and an inversion of the second allele, including a deletion of the start codon, resulting in a complete inactivation of Phafin2 (Figure S2).

### siRNA-mediated depletion of Phafin2

siRNAs against Phafin2 were described before (Pedersen et al). The following sequences were used to deplete Phafin2: siRNA 1 TGGTCAACCTTTAACTATA; siRNA2: GAAGCAAATACTAGACGTA. For siRNA transfections, 50 nM of siRNA was transfected using RNAiMax reagent. Transfected cells were analyzed 72h after transfection; knockdown efficiency was verified by western blotting.

### Live cell imaging

All live-cell imaging was performed on a Deltavision OMX V4 microscope (GE Healthcare) equipped with three watercooled PCO.edge sCMOS cameras, a solid-state light source and a laser-based autofocus. Environmental control was provided by a heated stage and an objective heater (20-20 Technologies). Cells were imaged in Live Cell Imaging buffer (Invitrogen) supplemented with 20 mM glucose. To enhance the rate of macropinosome formation, cells were stimulated by adding HGF (50 ng/ml) directly before imaging. For imaging of Lifeact-SNAP, cells were stained with SiR647-SNAP (New England Biolabs) according to the manufacturers protocol. Images were deconvolved using softWoRx software and processed in Fiji ^37, 38^.

### Immunostaining and stimulation of macropinocytosis

siRNA treated cells grown on glass coverslips were stimulated with HGF (50ng/ml) for 15 minutes in full medium (DMEM-F12, 10% FCS, Pen/Strep), fixed with 3% formaldehyde (Polysciences, 18814) for 15 min on ice, and permeabilized with 0.05% saponin (Sigma-Aldrich, S7900) in PBS. Fixed cells were then stained with a mouse anti-Rabankyrin-5 antibody (Abnova, CatNo: H0005 1479-B01P) at room temperature for 1 h, washed in PBS/saponin, stained with a fluorescently labeled secondary antibody (donkey anti-mouse Alexa555, Molecular Probes CatNo.: A31570) for 1h, washed in PBS and mounted with Mowiol (Sigma-Aldrich) containing 2 μg/ml Hoechst 33342 (Thermo Fisher Scientific, H3570).

### Quantification of Rabankyrin-5 dots

Wide field images of siRNA transfected cells labeled for endogenous Rabankyrin-5 were acquired automatically by an Olympus ScanR high content microscopy system using a UPLSAPO 40x objective, and analyzed automatically by the ScanR software. After back ground subtraction, dots of Rabankyrin-5 larger than 30 pixels were segmented automatically by the ScanR software based on an intensity threshold, and the number of dots per cell was measured. The total number of cells was quantified automatically by detection of Hoechst nuclear stain. Identical settings were used for all conditions within an experiment. In total more than 2500 cells (from at least 200 images) were analyzed for each condition, from three independent experiments.

### Confocal fluorescence microscopy

Confocal images were obtained using LSM710 confocal microscope (Carl Zeiss) equipped with an Ar-laser multiline (458/488/514 nm), a DPSS-561 10 (561 nm), a laser diode 405–30 CW (405 nm), and an HeNe laser (633 nm). The objective used was a Plan-Apochromat 63×/1.40 oil DIC III (Carl Zeiss). Image processing was performed with basic software ZEN 2010 (Carl Zeiss) and Fiji software.

### dSTORM imaging

Cells expressing Phafin2-GFP were fixed in 4 % PFA and labelled with anti-GFP-Nanobody (Atto488-conjugated), anti-Filamin A antibody (CF568-conjugated secondary antibody) and Alexa647-Phalloidin. Gold fiducials (100 nm) were added for image registration. Imaging was performed in 100 mM Tris, 50 mM NaCl with an oxygen scavenging system (10% Glucose, 10 kU catalase, 0.5 kU glucose oxidase) and 100 mM MEA as reducing agent. Imaging was performed using a Deltavision OMX V4 microscope (GE Healthcare); localization, reconstruction and alignment was performed in softWoRx software; all further image processing was performed in Fiji.

### Measurement of Dextran uptake by flow cytometry

Fluid phase uptake was assayed by measuring fluorescent dextran uptake using flow cytometry.

Fluorescent dextran (10 kDA (Fig. 5.3) or 70 kDA (Fig. 5.1), Molecular probes) was added to cells stimulated with HGF and cells were incubated for 20 min at 37°C. Cells were then washed 5 times with pre-warmed medium, trypsinized and dextran fluorescence was measured by flow cytometry.

When measuring Phafin2 KO cells, wild-type cells stably expressing H2B-mCherry were mixed and seeded out together with KO cells to minimize well-to-well variability. During flow cytometry measurements, wild-type and KO cells were gated according to their mCherry fluorescence and the fluorescence of Alexa488-conjugated dextran was determined in the respective populations.

All flow cytometry measurements were performed using an LSRII flow cytometer equipped with 488 and 561 nm laser lines.

### Membrane scission assays

Membrane scission was assayed by measuring the fluorescence of the pH sensitive RFP Phuji in response to pH changes. Phuji localized in sealed vesicles does not respond to changes of the extracellular pH, whereas surface-exposed fluorophores are readily quenched. Cells stably expressing Phafin2-GFP were seeded in Mattec dishes and transfected with pDisplay-Phuji. Imaging was performed in Live Cell Imaging buffer (Invitrogen) supplemented with 20 mM glucose. After being placed on the microscope stage, a perfusion pencil (Autom8 Scientific) was placed directly on top of the cell using a micromanipulator. Short bursts of imaging buffer (140 mM NaCl, 2.5 mM KCl, 1.8 mM CaCl2, 1.0 mM MgCl2, 20 mM PIPES or HEPES) buffered to pH 5.5 and 7.5 with PIPES and HEPES, respectively, were alternatingly perfused in 5 s intervals using a gravity driven perfusion apparatus (Warner). All solutions were heated using an in-line heating element (Warner). Perfusion intervals were controlled by an Arduino microcontroller using PyFirmata and custom Python scripts (available at https://github.com/koschink/Arduino_Perfusion).

### Protein-lipid interaction assays

In vitro lipid binding actitivies of Phafin2 PH and FYVE were determined by protein-lipid overlay assays ^20^. Protein expression, purification and lipid-overlay assays were performed as described ^39^.

### Yeast two-hybrid assays

Phafin2 interaction partners were identified by both surveying the IntAct database and by a yeast two-hybrid assay performed by Hybrigenics. In the latter case, Phafin2 was screened against a human monocyte-derived library. Using this approach, Filamin A was identified as two-hybrid interaction partner of Phafin2.

The interaction of Filamin A and Phafin2 was further characterized by measuring their interaction using a LexA-based yeast two-hybrid system. Phafin2 was cloned into pLexA, whereas Filamin A was cloned into pGAD-GH2. Both bait and prey plasmids were co-transformed into to the yeast strain L40 (ATCC strain MYA-3332) and resulting clones were used to assay interactions by LacZ assays. The interaction surface of Phafin2 with Filamin A was further mapped by using pLexA plasmids encoding Phafin2 truncation mutants instead of full-length Phafin2. To map the interaction domain of Filamin A with Phafin2, we generated constructs expressing subdomains of the identified Filamin A fragment encoding the CH2 and the first IG repeat in pGAD-GH2; also, these constructs were tested against full length Phafin2. Interaction strength was assayed by liquid beta-Galactosidase assays.

### Image processing and data analysis

All live cell images were deconvolved using Softworx (GE Healthcare) prior to analysis and presentation. All further image analysis and measurement steps were performed in Fiji using custom Jython scripts. Macropinosomes were manually tracked and a region of their limiting membrane was marked as region of interest. Fluorescence intensity of a circular ROI (10 pixel diameter) surrounding the marked region was quantified in all image channels and measurements were exported for further analysis. All further processing steps were performed using Python scripts and the “Pandas” data analysis package. Individual tracks of macropinosomes were temporally aligned, their fluorescence intensity normalized and the mean value and the SEM/95 % confidence interval was calculated for each timepoint. and plotted for presentation using either Matplotlib or, after export, Graphpad Prism. Analysis scripts are available at https://github.com/koschink/Phafin2.

### Quantification and statistical analysis

Statistical analysis was performed using Graphpad Prism. Student’s t-test (two-sided) was used as a measure for statistical significance. Data are always presented as mean +/- SEM, p values smaller than 0.05 were considered to be statistically significant. Sample sizes and p values are reported in the figure legends.

In the case of flow cytometry based Dextran uptake assays, measurements were normalized by setting the control measurements to 1 and a one-sample t-test (to account for the lacking variability.

All data presented is derived from at least 3 independent experiments; live cell imaging data is generated from multiple cells per experiment.

### Code availability

All custom software is openly available at GitHub and is implemented using freely available software packages (Python; ImageJ)

### Data availability

The authors declare that the main data supporting the findings of this study are available within the paper and its supplementary information files. Additional data are available from the corresponding authors on request.

## Supplemental Figure legends

**S1:**
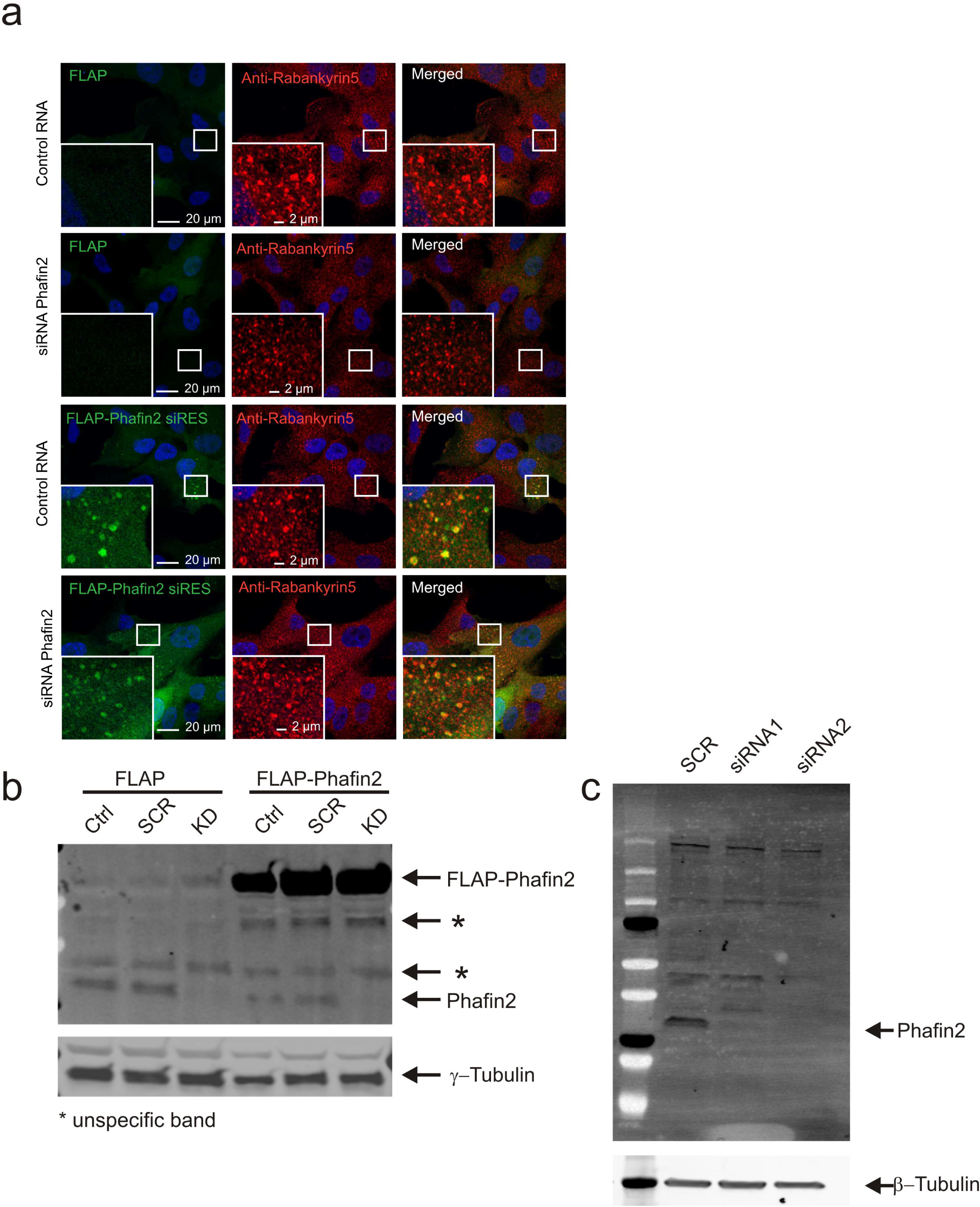
Depletion of Phafin2 impairs macropinosome maturation (related to Figure 5). a) Depletion of Phafin2 by siRNA results in smaller Rabanykrin5 vesicles. This can be rescued by expression of a siRNA-resistant Phafin2 construct. Shown are representative confocal images from experiment 5b. b) Representative Western blot showing knockdown efficiency and re-expression of FLAP-Phafin2 for one of the experiments in Figure. 5b. The other experiments showed similar knockdown efficiency (not shown). c) Representative Western blot showing knockdown efficiency for one of the experiments in Fig. 5a. The other experiments showed similar knockdown efficiency (not shown).

**S2:**
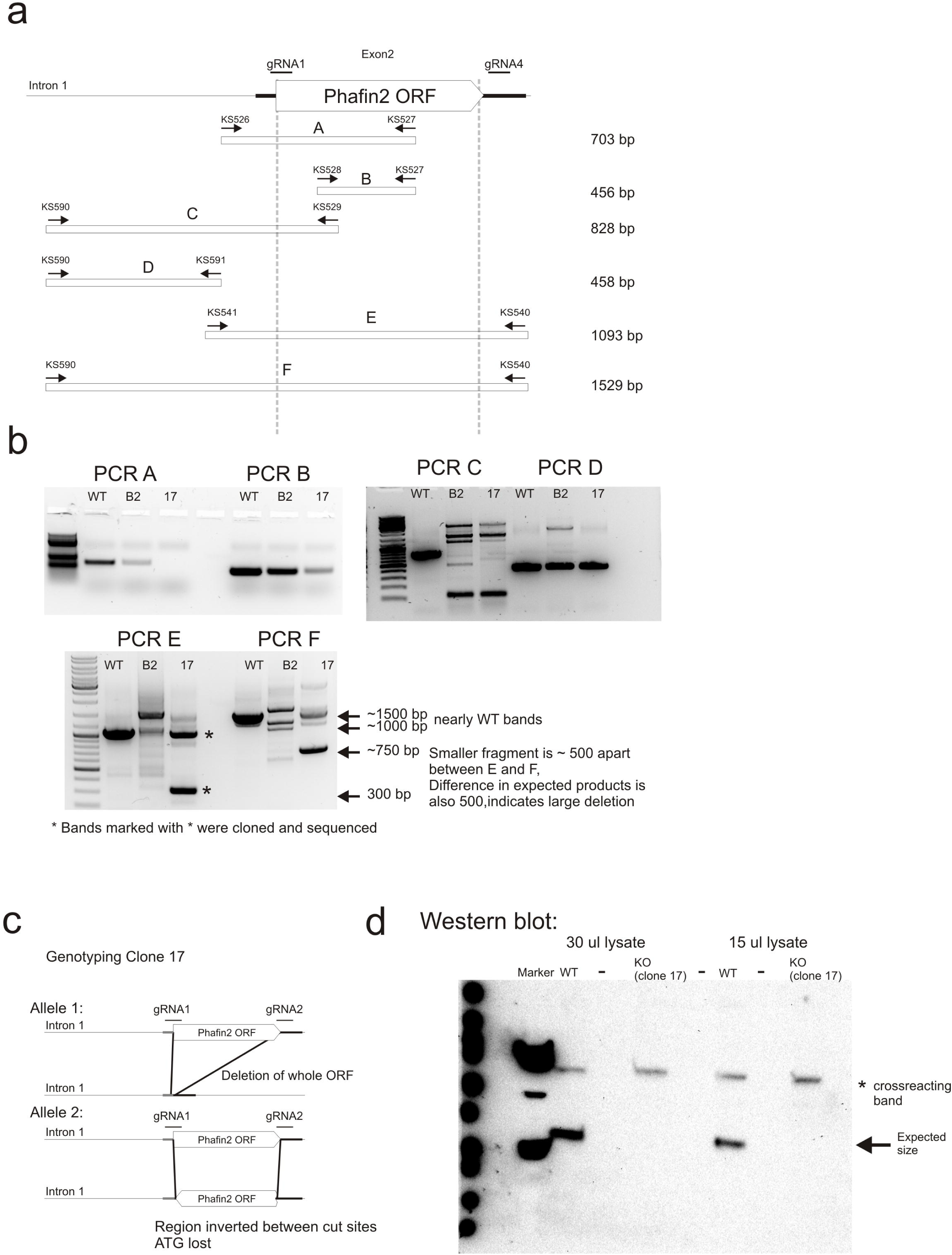
Characterization of CRISPR-generated Phafin2 KO cell lines (related to Figure 5). a) Schematic display of the Intron-Exon structure of Phafin2, localization of gRNAs, Primer combinations and expected product sizes used to characterize Phafin2 KO cell lines. b) PCR products from Phafin2 KO cell lines. c) Genomic organization of the used Phafin2 -/- cell line. d) Western blot showing complete absence of Phafin2 in the used Phafin2 -/- cell line.

**S3:**
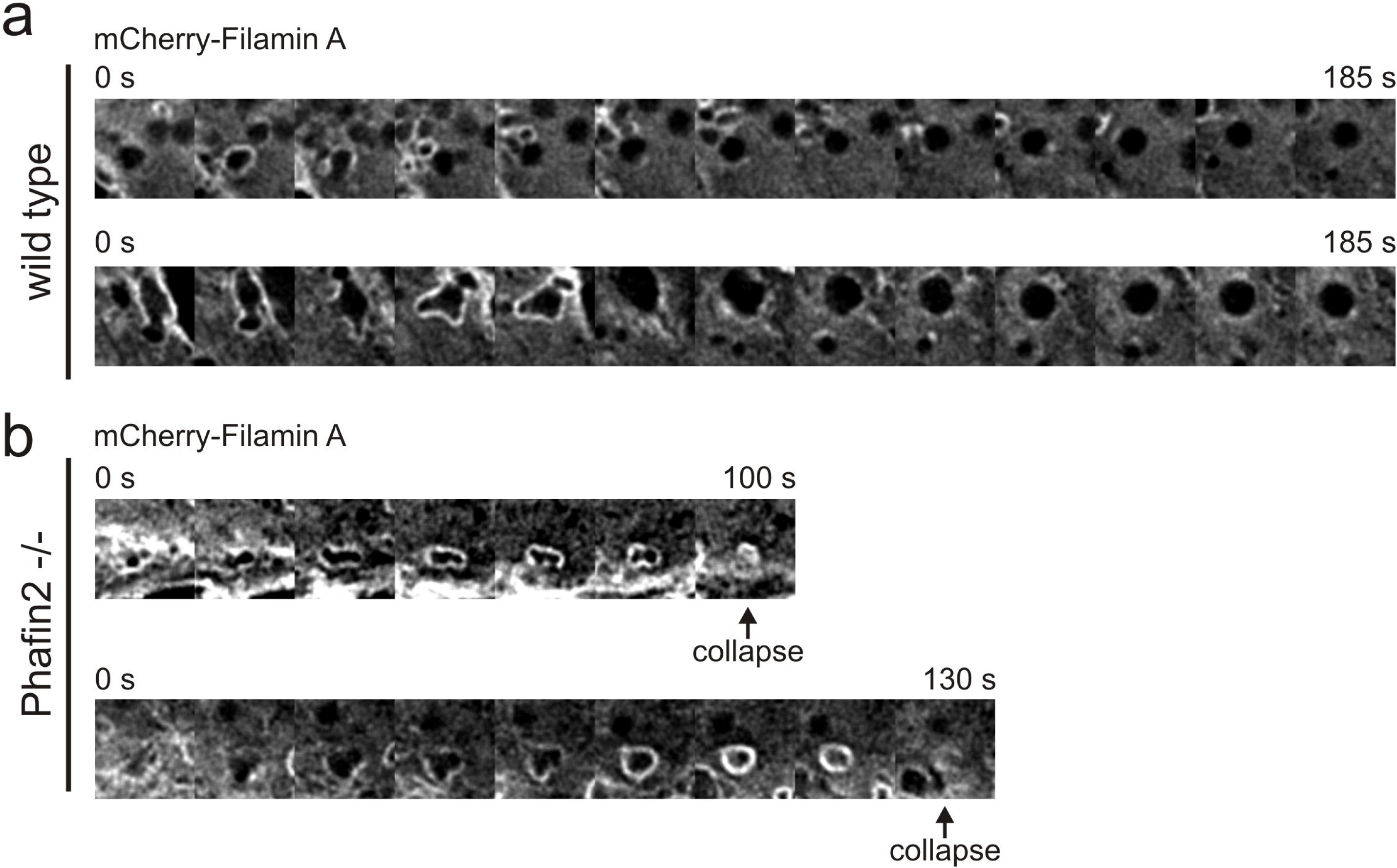
Deletion of Phafin2 impairs Filamin A removal from macropinosomes (Related to Figure 7). a) wild-type cells show transient Filamin A recruitment to newly-formed macropinosomes; as Filamin A is removed, macropinosomes mature to endosomal stages. b) Macropinosomes in cells lacking Phafin2 retain Filamin A and frequently collapse. Shown are representative time series from Figure 7.

**Supplemental Figure S4:**
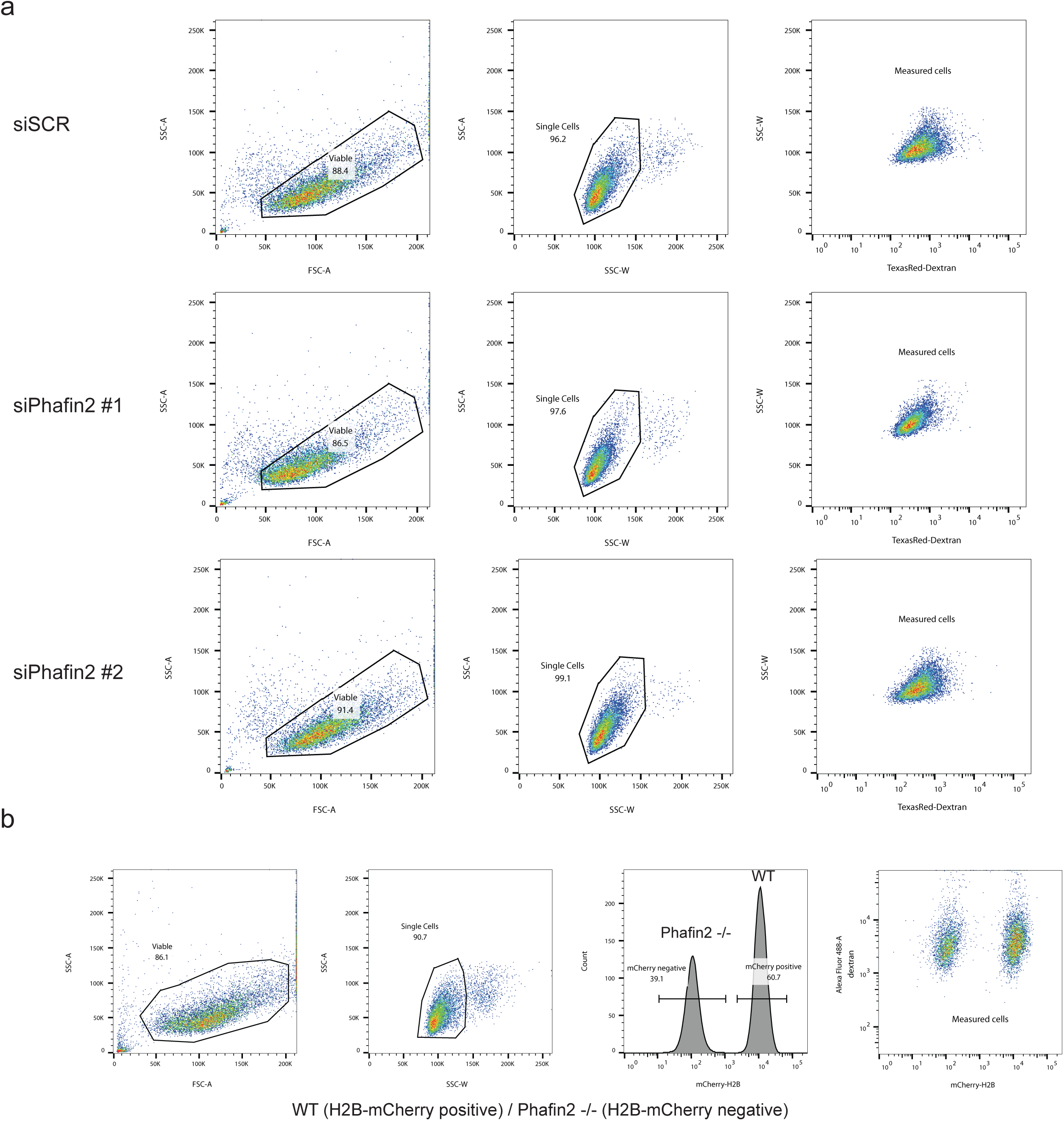

**Table S1:**
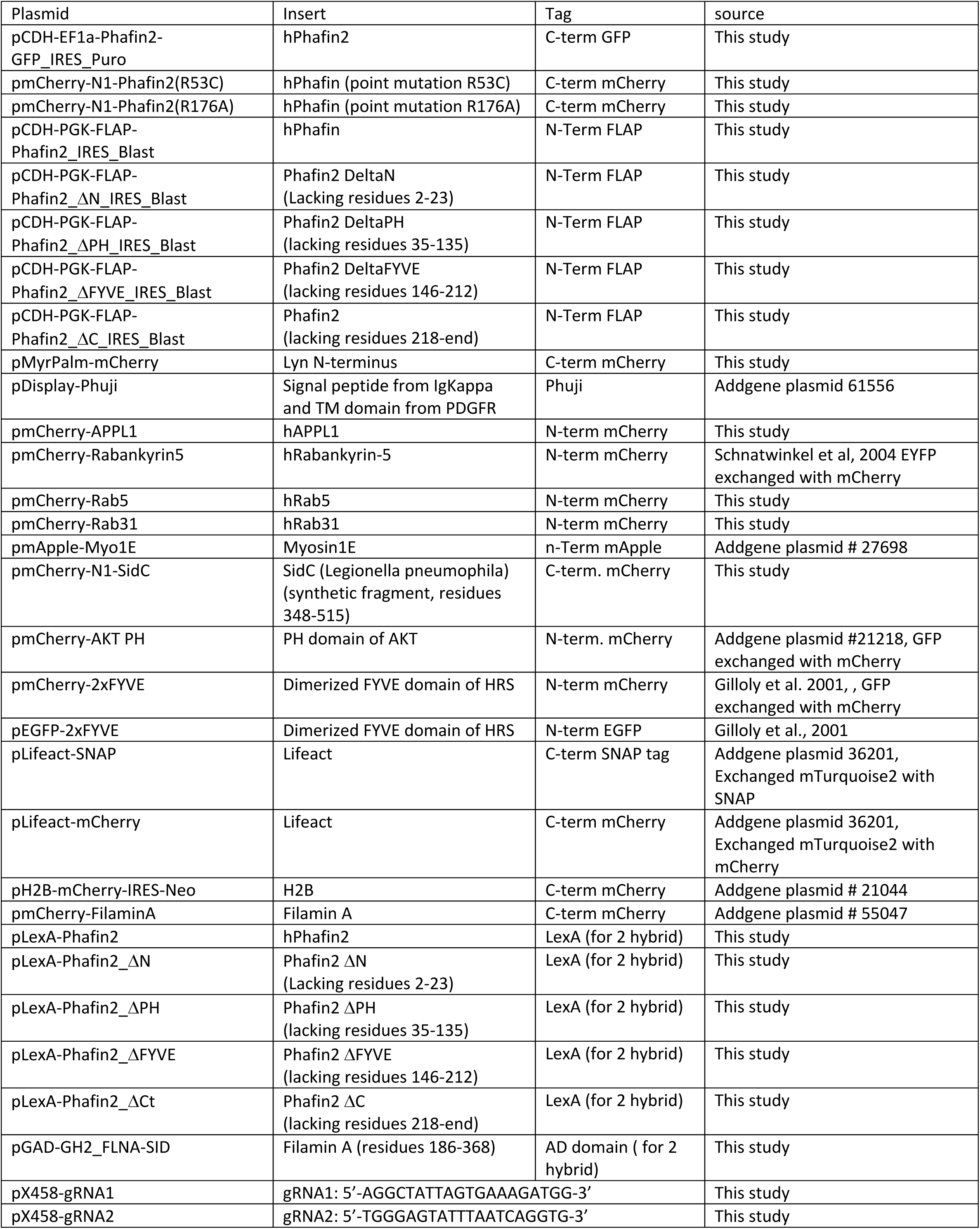
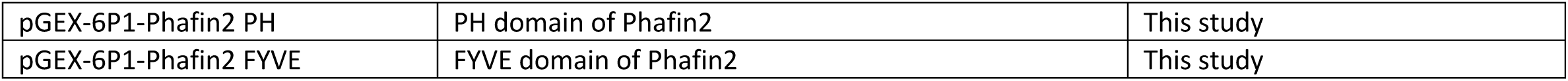
Plasmids used in this study

## Supplemental movie legends

S1: Phafin2-GFP localizes to macropinosomes directly after cup closure, indicated by the plasma membrane probe MyrPalm-mCherry (related to Figure 1).

S2: Phafin2 localizes to macropinosomes after scission (related to Figure1). Cells expressing surface-displayed pH sensitive RFP (Phuji) and Phafin2-GFP were perfused with pH 5.5 and 7.5, leading to quenching and unquenching of Phuji fluorescence. Phafin2 is localized to sealed vesicles.

S3: The first localization of Phafin2 occurs prior to recruitment of the early endosomal protein APPL1 (related to Figure 2).

S4: A functional Phafin2 FYVE domain is required for membrane association (related to Figure 3). Wild-type Phafin2 shows a biphasic localization, whereas a FYVE domain mutant shows a completely cytosolic localization.

S5: A functional PH domain is required for localization to newly-formed macropinosomes (related to Figure 3). While wild-type Phafin2 shows a biphasic localization, a mutant with an inactive PH domain does not show the first localization phase on newly-formed macropinosomes, but is still able to localize to the second, endosomal phase.

S6: The first phase of Phafin2 localization is independent of Vps34 activity (related to Figure 4). Inhibition of Vps34 does not affect the first transient localization phase of Phafin2 to nascent macropinosomes, but completely abolishes the second localization phase to endosomal stages.

S7: Macropinosome formation in wild-type and Phafin2 KO cells (related to Figure 5). Wild-type cells show robust formation of new macropinosomes, as indicated by formation of large, 2xFYVE-positive vesicles; Phafin2 KO cells only rarely form large macropinosomes, but rather show formation of multiple small vesicles.

S8: Newly-formed macropinosomes are surrounded by actin and are subjected to actin-based forces related to Figure 6).

S9: Phafin2 shows transient colocalization with Filamin A on newly-formed macropinosomes, but not on endosomal stages (related to Figure 6).

S10: Newly-formed macropinosomes enter the cell through gaps in the surrounding Filamin A network (related to Figure 6).

S11: Effect of Phafin2 knockout on Filamin A recruitment (related to Figure 7). Wild-type cells show transient Filamin A localization to newly formed macropinosomes; Filamin A is shed from macropinosomes during their maturation. Macropinosomes in Phafin2 KO cells show prolonged Filamin A localization and are unable to shed Filamin A, leading to frequent back-fusion.

